# Gene-scale *in vitro* reconstitution reveals histone acetylation directly controls chromatin architecture

**DOI:** 10.1101/2024.11.08.622658

**Authors:** Yohsuke T. Fukai, Tomoya Kujirai, Masatoshi Wakamori, Setsuko Kanamura, Lisa Yamauchi, Somayeh Zeraati, Chiharu Tanegashima, Mitsutaka Kadota, Hitoshi Kurumizaka, Takashi Umehara, Kyogo Kawaguchi

**Affiliations:** Nonequilibrium Physics of Living Matter RIKEN Hakubi Research Team, RIKEN Center for Biosystems Dynamics Research, 2-2-3 Minatojima-minamimachi, Chuo-ku, Kobe 650-0047, Japan; Laboratory of Chromatin Structure and Function, Institute for Quantitative Biosciences, 1-1-1 Yayoi, Bunkyo-ku, Tokyo 113-0033, Japan; Laboratory for Epigenetics Drug Discovery, RIKEN Center for Biosystems Dynamics Research, 2-2-3 Minatojima-minamimachi, Chuo-ku, Kobe 650-0047, Japan; Laboratory for Developmental Genome System, RIKEN Center for Biosystems Dynamics Research, 2-2-3 Minatojima-minamimachi, Chuo-ku, Kobe 650-0047, Japan; RIKEN Cluster for Pioneering Research, 2-2-3 Minatojima-minamimachi, Chuo-ku, Kobe 650-0047, Japan; Institute for Physics of Intelligence, Department of Physics, The University of Tokyo, 7-3-1 Hongo, Bunkyo-ku, Tokyo 113-0033, Japan

## Abstract

Chromatin organization plays a crucial role in gene regulation [1, 2, 3], but disentangling the contributions of various epigenetic components to gene-scale chromatin structure remains challenging. While *in vitro* chromatin reconstitution enables controlled studies on the effect of biochemical factors on the structure, current methods are either limited to short arrays or lack control over histone modification patterns. Here we directly test how histone modification affects higher-order chromatin architecture by characterizing gene-scale reconstituted chromatin using single-molecule microscopy and *in vitro* Hi-C. We reconstitute 20-kilobase chromatin arrays with histone modification patterns controlled at 12-nucleosome resolution, achieving complete assembly of 96 nucleosomes in the designed order as confirmed by atomic force microscopy and longread sequencing. Observing end-to-end fluctuations of the reconstituted arrays, we find that increasing the density of acetylated nucleosomes leads to larger structural fluctuations with longer relaxation times, consistent with the predictions of a polymer model with hydrodynamic interactions. We demonstrate through *in vitro* Hi-C how acetylation reduces contact frequency between nucleosomes and induces open conformations. In heterogeneously modified arrays, differential contact probabilities between acetylated and unmodified regions lead to distinct structural domains. The results establish the physical principles by which histone modifications directly modulate chromatin architecture through altered nucleosome-nucleosome interactions, providing a quantitative framework for understanding and engineering genome organization.

---

The dynamic regulation of chromatin structure plays a crucial role in establishing and maintaining cell identity of eukaryotes [1, 2, 3]. Distinct histone modification patterns have been correlated with three-dimensional genome structure [4, 5, 6]. However, the molecular mechanisms underlying these structures are multifaceted; they encompass not only direct physical interactions between nucleosomes [7, 8, 9] but also the action of various proteins that recognize specific epigenetic modifications [10, 11, 12, 13, 14]. The limited control over the chromatin environment *in vivo* makes it challenging to disentangle the contributions and interdependencies of diverse mechanisms shaping large-scale chromatin structure and dynamics across genes or regulatory domains.

Reconstituted chromatin systems have been a promising approach to overcome these limitations, offering a controllable platform for dissecting the impact of specific factors starting at the single nucleosome level [9, 14, 15]. Experiments using reconstituted regularly-spaced arrays of 12 nucleosomes (a 12-mer) have demonstrated that the structure depends on the acetylation level of histones [7, 8], binding to proteins [16, 17] and linker length [18], and can also undergo liquid-liquid phase separation or aggregation at high concentrations [19, 20]. However, the complex, large-scale organization of chromatin *in vivo*, particularly within and beyond gene-sized domains (> 10 kb) with heterogeneous histone modifications [21], is difficult to reproduce *in vitro*. Recreating these complex, patterned structures is crucial for unraveling the principles governing the large-scale chromatin architecture, including the impact of long-range interactions beyond tens of nucleosomes.

To address the complexity of large-scale chromatin organization, we developed a method to reconstitute long chromatin with defined histone modification patterns. By constructing a 20-kb, 96-mer nucleosome array with histone modification patterns controlled at 12-mer resolution, we uncover the relationship between histone modification patterns, structure, and dynamics at the scale involving distal interactions within the chromatin chain. Utilizing the barcodes introduced at single nucleosome resolution, we further probe the conformation of the reconstituted nucleosome arrays through *in vitro* conformation capture experiments (i.e., *in vitro* Hi-C). The obtained contact map validates how the patterning of histone modifications is sufficient to induce heterogeneous contacts at kilobase scales, resembling higher-order structures observed *in vivo*, such as TADs and compartments.

## Reconstituting chromatin with defined nucleosome modifications

To construct long arrays of nucleosomes with defined histone modification patterns, we first reconstituted 12-mer nucleosome arrays with distinctly modified human histone octamers and subsequently pooled them to ligate up to 96-mers (Fig. 1a). A similar strategy has been taken to make shorter arrays (four to 25-mers) [22, 23, 24]. We designed the DNA of 12-mer arrays consisting of a repeat of the Widom 601 sequence [25] with distinct sticky ends that can only be ligated in the designed order (Fig. 1b), allowing the creation of defined sequences of histone modification patterns at 12-mer resolution. We chose histone tail acetylation in this study considering its well-established role in gene activation and its demonstrated capacity to directly influence chromatin structure. We used H4 tail tetra-acetylation (i.e., H4K5K8K12K16ac) to reproduce the H4 hyperacetylation found in the transcriptionally active chromatin [26, 27, 28, 29]. The acetylated 12-mers were combined with unmodified 12-mers to construct four distinctly patterned arrays: all acetylated (All-Kac), all non-acetylated (All-nonKac), center region acetylated (Center48-Kac), and 12-mer alternatingly acetylated (Every12-Kac) patterns (Fig. 1c and Extended Data Fig. 1a).

**Fig. 1:**
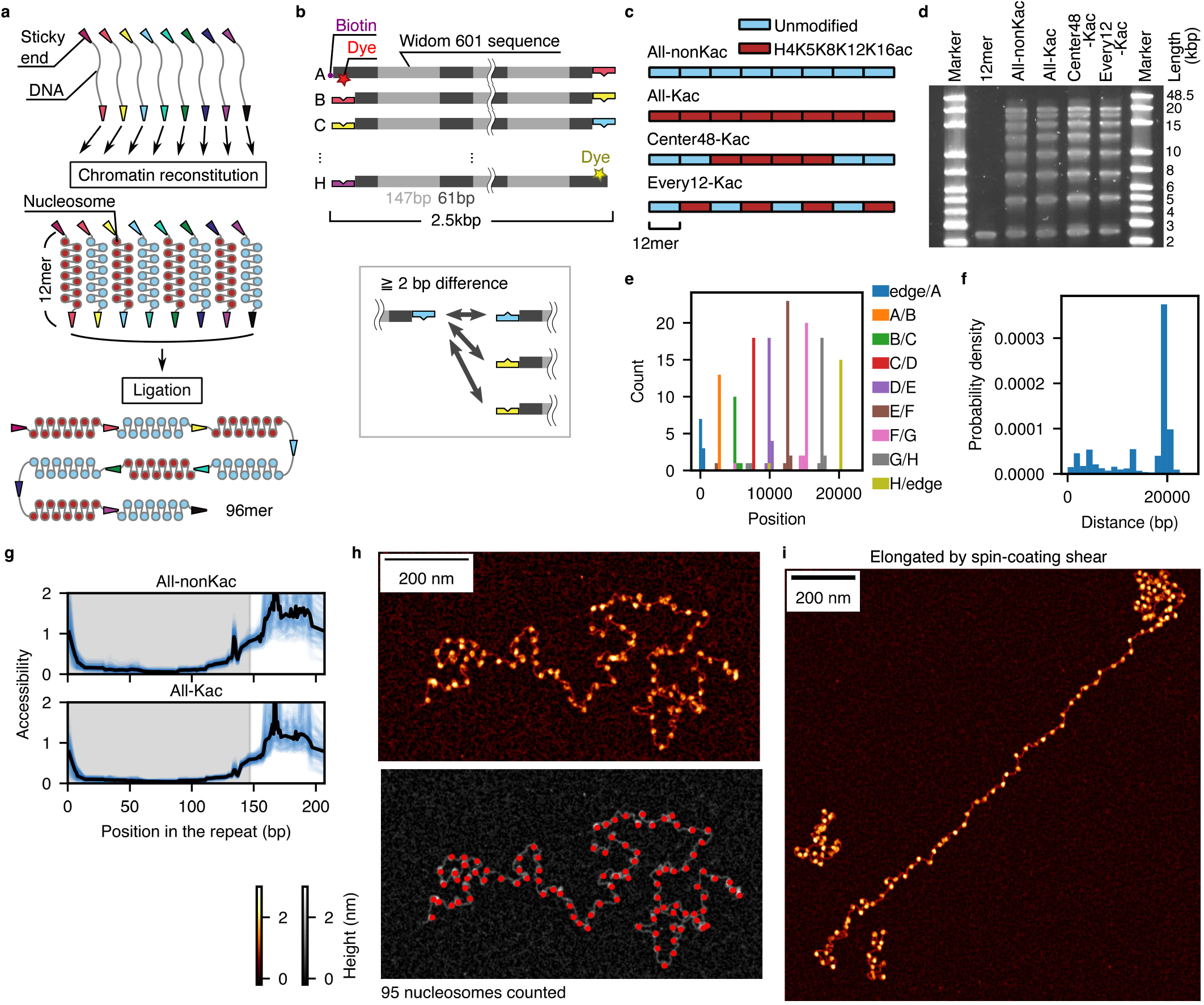
Reconstituting chromatin arrays with defined histone modifications. **a**. Schematics of the experiment. **b**. Design of the eight 12-mer arrays. **c**. Design of the histone modification patterns. **d**. Electrophoresis result of the ligated product of eight 12-mer arrays. **e**. Counts of the ligated regions at each position found in Nanopore sequences. **f**. Sequence bias-corrected distribution of the length of non-barcoded chromatin when subsampling the sequences with both ends of the 96-mer design. **g**. Accessibility of the DNA in all repeats in the long chromatin arrays. The gray-shaded region indicates the 601 sequence, and the white region indicates the linker. **h**. AFM image of a reconstituted long array. Each observed blob likely corresponds to a nucleosome, resulting in a total count of 95 nucleosomes. The red points show the detected nucleosome positions. **i**. AFM image of a reconstituted 96-mer chromatin array observed after applying shear by the spin coater to elongate the chains (see Methods). The image also shows two shorter arrays, identified as 12-mer and 24-mer arrays.

In order to construct long nucleosome arrays with high histone-loading fractions on 601 sequences, we first reconstituted the 12-mer arrays with an optimized method (see Methods). Gel shift assays confirmed efficient reconstitution of the 12-mers, showing that less than 5% of the DNA remained unbound to histones (Extended Data Fig. 1b-i). We also checked the uniformity of the reconstitution visually by atomic force microscopy (AFM); around 90% of detected arrays counted 12 blobs that correspond to nucleosomes (Extended Data Fig. 1j,k).

To efficiently ligate the 12-mers into larger structures, we optimized the concentration of the samples, magnesium ion, and adenosine triphosphate (ATP); magnesium ions induce undesired aggregation for nucleosomes but are necessary for the ligation reaction by T4 ligase. With the ligation condition that yielded 96-mers with a high fraction (Fig. 1d), the nucleosomes remained intact after the ligation as assessed by the gel shift assay (Extended Data Fig. 2a). By Nanopore sequencing, we also confirmed that the ligation sites identified in the long reads corresponded to the designed positions (Fig. 1e and Extended Data Figs. 2b), and the length distribution of strands containing both end sequences had a sharp peak around 20 kb, matching the design (Fig. 1f and Extended Data Fig. 2c). The positioning of the histones on the 601 sequence positions was also confirmed by the accessibility assay by adenine methylation [30, 31] (Extended Data Fig. 3), where enzyme access was found to be almost completely suppressed at the positions of histones (Fig. 1g).

We further validated the ligated products by AFM (Fig. 1h,i and Extended Data Fig. 4). By optimizing the imaging protocol, we were able to reach a resolution where we could count the number of histones contained in the nucleosomes, which were close to 96 for the largest structures (Fig. 1h,i and Extended Data Fig. 4a-f). In the AFM images, we also observed intermediate products corresponding to the heterogeneous population of arrays in multiples of 12-mers (Fig. 1d,i and Extended Data Fig. 4g-m).

### Spatio-temporal fluctuation of nucleosomes depends on the fraction of acetylation

Next, we aimed to characterize the spatio-temporal fluctuation of 96-mers and its dependence on histone modification patterns by single-molecule microscopy. Fully ligated arrays should contain distinctly colored fluorescent probes at both ends (Figs. 1b and 2a). We designed an observation chamber passivated with polyethylene glycol to minimize non-specific binding to the substrate (see Methods). The chamber can introduce fluid flow, which allows the exchange of the buffer into distinct salt conditions, as well as the application of shear force to check the extent of elongation of the molecules (Fig. 2b,c). In our observation of single molecules, we found non-specific binding of the nucleosomes to the substrate, likely due to defects in the passivated surface of the glass. We excluded non-specifically bound molecules post-imaging, and focused on molecules that fluctuated significantly during the observation and elongated sufficiently upon the application of shear (Extended Data Fig. 5a-c).

**Fig. 2:**
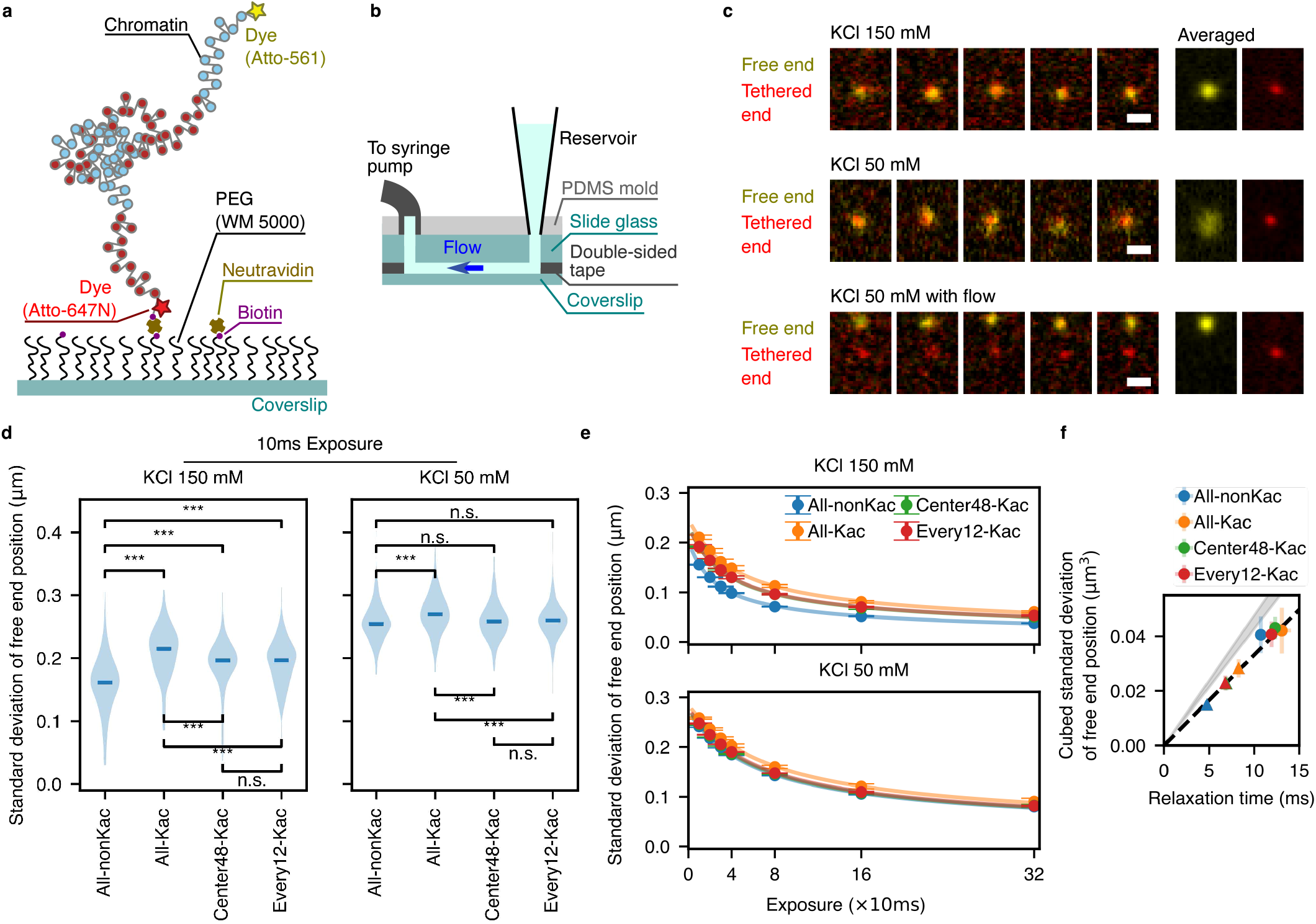
Single molecule microscopy shows histone modification-dependent fluctuations. **a**. Schematic of the single-molecule observation of a 96-mer array. **b**. Design of the chamber. **c**. Montage of the fluorescent probe observations tagged at the free and tethered ends of the chromatin molecules. The scale bar is 1 µm. **d**. Extent of fluctuation for a single frame (10 ms exposure) measured from the fluctuation amplitude of the free ends. The asterisks indicate the statistical significance of the Mann-Whitney test adjusted by the Benjaminini-Yekutieli method: “***” corresponds to statistical significance with *p* < 0.001. “n.s.” means not significant. **e**. The median value of the extent of fluctuation with varied exposure time and the theoretical fits. The error bars represent the bootstrap standard deviations for the median value estimation. **f**. Spatial extent of fluctuation in the zero-exposure limit plotted with respect to the relaxation time. Circle: KCl 50 mM, triangle: KCl 150 mM. The error bars represent the combined bootstrap standard deviations of two replicates. The grey shaded area represents the Zimm model prediction for the tethered chain setup, using the viscosity of water between 25 and 30 °C.

The molecules presented fluctuation at the size of around 100 − 200 nm (Fig. 2c,d and Extended Data Fig. 5d), and elongated to the length of 750 1500 nm when applied shear, consistent with the expected size of 96-mer nucleosomes (Extended Data Fig. 5b,c). The amplitude of the fluctuation had a clear histone modification dependence: acetylated nucleosomes had larger fluctuation compared with the non-acetylated nucleosomes, consistent with the acetylated histones having weaker attractive interactions between the histone tails and negatively charged regions such as the acidic patches of the nucleosomes and DNA. Interestingly, this difference was enhanced in the physiological 150mM salt condition and was not seen for 50 mM, indicating that the charge difference induced by acetylation is a subtle effect that depends significantly on the environment. At lower salt concentrations, repulsive interactions between DNA regions can dominate, whereas at higher salt concentrations, electrostatic screening mitigates these DNA-DNA repulsions, allowing histone-histone and/or histone-DNA interactions to become prominent. We also found that the patterned chromatin with half of the histones being acetylated (Center48-Kac and Every12-Kac) tended to fluctuate in size between the fully-acetylated and non-acetylated arrays (Fig. 2d and Extended Data Fig. 5d).

We found that the size of molecules observed by AFM had a similar trend; acetylated arrays tended to be larger compared to the non-acetylated, a difference undetectable at the 12-mer scale but becoming significant beyond 24-mers (Extended Data Fig. 6a-c). The diffusion constant of 12-mer nucleosomes measured for free-floating single molecules did not significantly differ between acetylated and non-acetylated arrays (Extended Data Fig. 7a). Previous studies have reported ≈ 20% changes in the sedimentation coefficient of 12-mer nucleosomes upon acetylation [7, 8], highlighting that acetylation-dependent effects for 12-mers may only become apparent under the influence of an external force such as during centrifugation. The difference between the distinct patterns of acetylation, the Center48-Kac and Every12-Kac arrays, was not detected in the single-molecule fluorescence signal (Fig. 2d and Extended Data Fig. 5d) and AFM (Extended Data Fig. 6d), indicating that the pattern dependence in heterogeneously modified chromatin is small if the overall acetylation fraction is the same.

From the live imaging data, we further estimated the true width of the end-to-end fluctuation as well as its time scale of relaxation. This was achieved by changing the number of averaging frames to calculate its effect on the fluctuation amplitude and fitting it with a theoretical model accounting for the finite camera exposure time (Fig. 2e and Extended Data Fig. 5e, see Methods). The time scale of relaxation, around ten milliseconds or less (Fig. 2f), is shorter than the time scale found in the longest lifetime of stacked nucleosomes [32], which was reported to be in the order of 100 ms [17]. We also observed that the time scale of relaxation was slower for chromatin arrays containing acetylation. Altogether, these results suggest that constructed 96-mer nucleosomes were fluctuating freely in a histone mark density-dependent manner, without forming aggregates.

### Physical model explains the relationship between fluctuation amplitude and relaxation time scale

The relaxation time scales of the chromatin fluctuations were proportional to the cube of their spatial extent (Fig. 2f). This cubic relation suggests that the dynamics of our reconstituted chromatin is consistent with the Zimm model [33, 34], which accounts for hydrodynamic interactions in polymer dynamics. The Zimm model predicts that the slowest relaxation time *τ* follows *τ* ∝ *ηR*^3^/*k*_B_*T*, where *R* is the root-mean-squared end-to-end distance, *η* is the solvent viscosity, *k*_B_ is the Boltzmann constant, and *T* is the temperature. As an independent measurement, we observed that the diffusion of 12-mer nucleosomes was faster than 1/12 of the mononucleosome diffusion constant, aligning closely with the Zimm model’s prediction of inverse square-root scaling (Extended Data Fig. 7a) [35].

For 25 to 30 °C water, the Zimm model for a tethered chain predicts *R*^3^/*τ* ≈ 4.27 to 4.77 µm^3^/s. Although there is ambiguity in the prefactor in this theory (see Supplementary Methods, Extended Data Fig. 7b,c), the order matches with the slope yielded from experimental data, 3.3 ± 0.1 µm^3^/s (Fig. 2f). Several factors may be combined to explain the difference between theory and experiment, such as the deviation from a beads-on-a-spring type of model, as well as increased viscosity near the coverslip surface [36].

We can consider that the spatial extent of chromatin fluctuation, *R*, depends on the Kuhn length *d*, which is influenced by ionic conditions and histone mo<u>di</u>fications. Assuming a Gaussian chain, 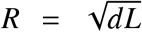, where *L* is the contour length. For *L* = 2.0 µm, observed *R* values of 250 nm (All-nonKac) and 300 nm (All-Kac), both at 150mM salt, correspond to Kuhn lengths of 31 nm and 47 nm, respectively. With a linker DNA length of 21 nm, the increased Kuhn length upon acetylation suggests a relatively stronger repulsive interaction between next-nearest neighbor nucleosomes (separated by two linkers), resulting in a stiffer chain.

### Chromatin contact pattern depends on the histone modification pattern

To further probe the structural changes associated with histone modification patterns, we reconstituted nucleosome arrays using DNA with linker-barcoded arrays to measure the contact frequency by the sequencing-based chromosome conformation capture method [37], which we call *in vitro* Hi-C (Fig. 3a,b). We first confirmed that the basic properties of the nucleosome arrays such as nucleosome occupancy, linker accessibility, and ligation order did not change due to the barcodes (Extended Data Figs. 1d-i,2d-j). Next, we optimized the fixation, digestion, and ligating steps for the barcoded 601 array from the method proposed for *in vitro* reconstituted yeast chromatin [38] (Supplementary Methods, Extended Data Fig. 8). The optimized protocol obtains a high yield of contacts between non-adjacent regions within the same molecule while suppressing nonspecific contacts quantified by the intermolecular contacts (Fig. 3c and Extended Data Fig. 9a, Methods).

**Fig. 3:**
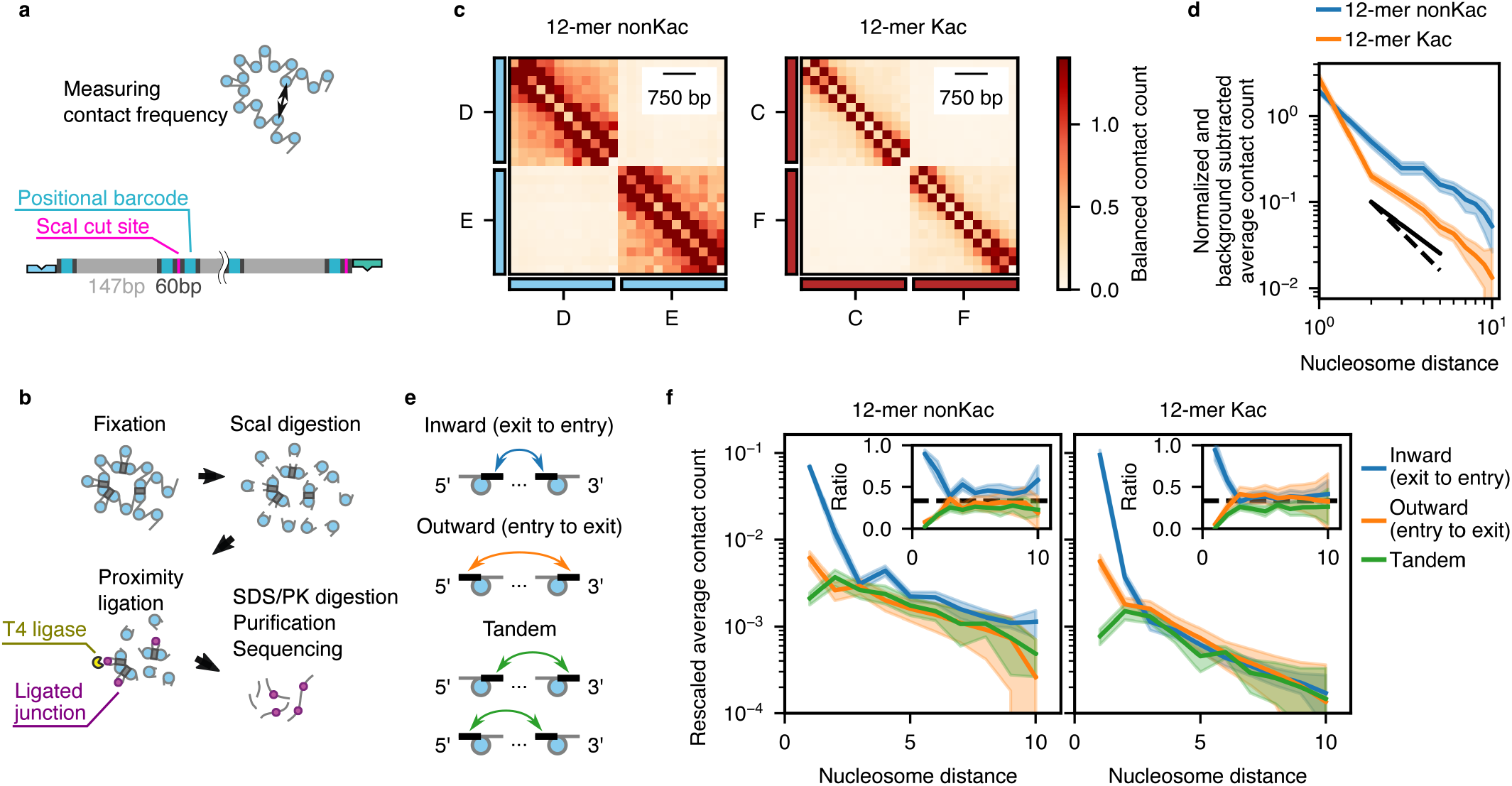
*In vitro* contact maps of reconstituted 12-mers. **a**. Schematic of the experiment and design of the barcoded 12-mer arrays. **b**. Illustration of the protocol. **c**. Contact map obtained from the mixed 12-mers with distinct barcodes. The contact is balanced and normalized so that the average contact frequency is 1. **d**. Contact frequency as a function of the nucleosome position. The shaded area represents the uncertainty (twice the s.e.m.). The dashed lines indicate the average contact frequency of the barcodes from the different 12-mer arrays. The black solid and dashed lines are the guide for the eye with the exponent − 1.5 and − 2, respectively. **e**. Schematic of the contact orientation. **f**. Contact frequency classified by orientation as a function of the nucleosome position distance. The shaded area represents uncertainty (twice the s.e.m.). The inset shows the ratio to the total contact frequency. The dashed lines are at ratio = 1/3.

For both the non-acetylated and acetylated 12-mer arrays, we found power-law scaling in terms of the distance versus contact frequency, which is consistent with a Gaussian chain model (Fig. 3d and Extended Data Fig. 9b). Interestingly, however, the decay of contact frequency occurred from a shorter distance in the acetylated sample compared with the non-acetylated arrays, resulting in a significantly decreased number of contacts for longer distances in the acetylated arrays (see Methods).

Utilizing the single-nucleosome resolution of the barcodes, we further dissected the contact frequency in the different orientations (Fig. 3e), inspired by the method employed in Ref. [39]. The result shows that the orientation dependence quickly decays and becomes indistinguishable beyond around the 5-mer distance (Fig. 3f and Extended Data Fig. 9c,d), suggesting that there is no canonical structure in our 12-mer samples [40], such as 30 nm fibers that have been observed for arrays with linker histone proteins or shorter linker length [41, 42, 18, 43, 44]. This result is consistent with there being no evidence of stacked structure observed in our AFM observation, as well as in the time scale of single-molecule fluctuations.

Next, we conducted the conformation capture experiment using the longer ligated chromatin. By this method, we detected contacts up to distances of more than 60-mers (around 12 kb) at significant levels beyond the inter-array contacts. The contact map of the heterogeneous chromatins, the Every12-Kac arrays and Center48-Kac arrays, were clearly distinct (Fig. 4a). This difference in the structure can be explained by the reduced contact probability between the acetylated pairs compared with the unmodified pairs (Fig. 4b). Compared with the uniformly patterned nucleosomes, the fluctuations in the contact frequency of the proximal 12-mers were larger in the Center48-Kac arrays (Fig. 4c and Extended Data Fig. 10a), reflecting the inhomogeneity of intramolecular contacts existing specifically in the Center48-Kac arrays. Consistent with these, the contact frequency between the nonKac regions was the highest, whereas the contact between Kac regions was less frequent (Fig. 4d and Extended Data Fig. 10b-d). These results indicate that the heterogeneous histone modification patterns on the chromatin array are enough to induce TAD-like contact patterns [1].

**Fig. 4:**
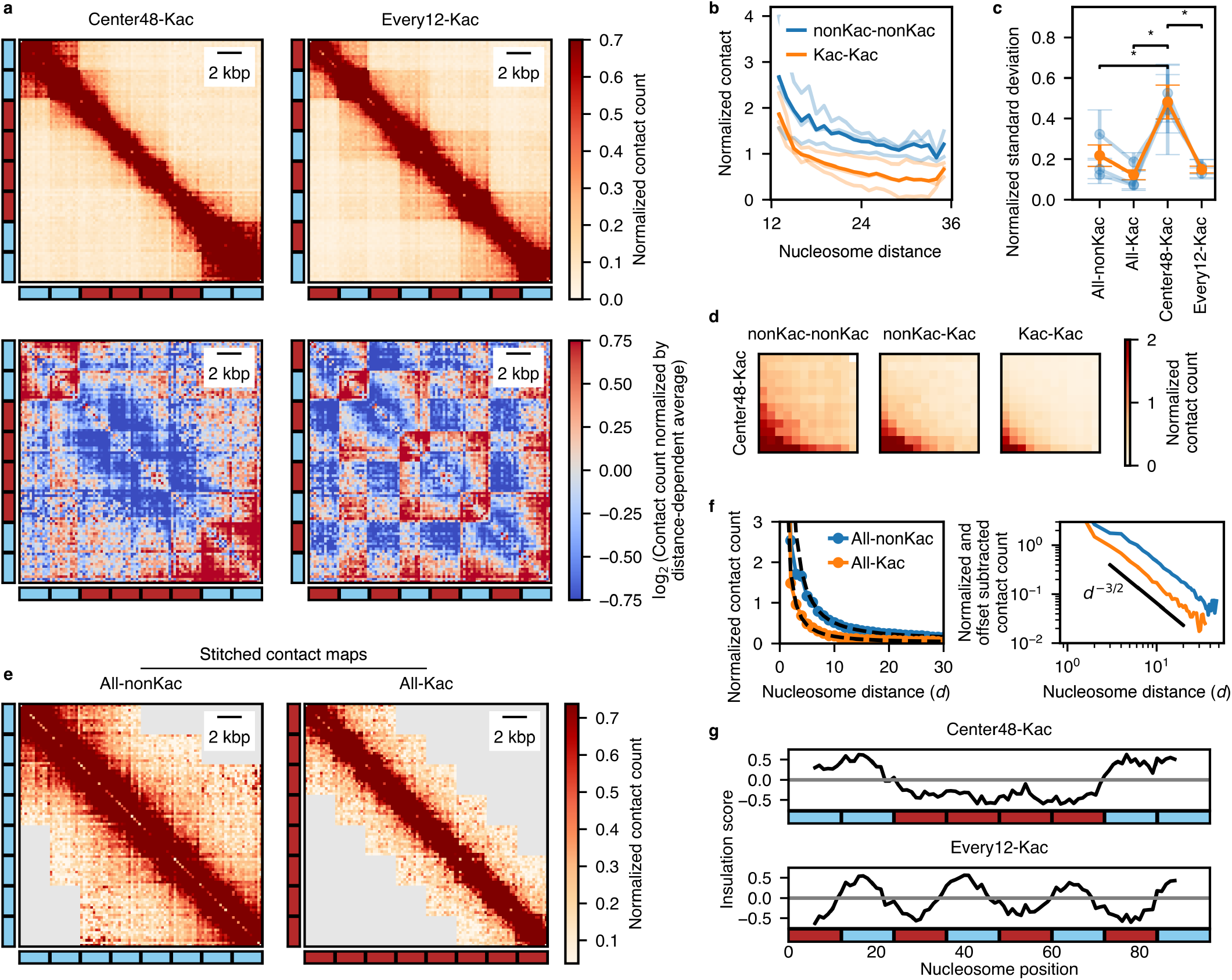
*In vitro* contact maps of reconstituted chromatin with heterogeneous histone modifications. **a**. Contact map of the ligated nucleosome arrays (96-mers as the largest) with inhomogeneous patterns. (top) Raw contact counts normalized to the total count. (bottom) The logarithm of counts normalized by the distance-dependent average. **b**. Average number of contacts between the second adjacent 12-mer regions for the Every12-Kac arrays as a function of the nucleosome distance. The light-colored lines are the replicates, while the dark-colored lines are the average. The total average values were significantly different by *p* < 10^−4^ in the Mann-Whitney test for all individual replicate pairs as well as their average. **c**. Standard deviation of the median contact between the adjacent 12-mer regions, rescaled by the mean contact frequency of all pairs. “*” indicates the statistical significance in the Brown-Forsythe test with *p* < 0.05. The light blue symbols are replicates, while the solid orange symbols are the average, both with s.d. as the error bars. **d**. Accumulated contact frequency between adjacent 12-mer regions for the Center48-Kac arrays. **e**. Stitched contact map of the homogeneous arrays. The gray area shows regions with non-significant contact counts in which more than 10% of the bins have negative contacts after background subtraction. **f**. (top) Contact frequency as a function of the nucleosome position distance in the stitched contact map. The dashed lines are the power-law fits with the exponent − 1.5 with offset. (bottom) The offset-subtracted contact frequency as a function of the nucleosome position distance. The black solid line is the guide for the eye with the exponent − 1.5. **g**. Insulation score for the stitched inhomogeneous arrays with a window size of 4 as a function of the nucleosome position.

For the uniformly patterned arrays (Extended Data Fig. 10e), we asked how the contact frequency decays for longer distances than 12-mers and how the decay depends on acetylation. We reconstructed the contact map at the 96-mer scale (Fig. 4e), assuming that the polymer structure is translationally symmetric and also that the contact map of subregions will be the same irrespective of it being probed in a large or small array (see Methods). Calculating the distance dependence of the contact frequency, we again obtained consistent results with a Gaussian chain, where the frequency decays with the power close to −1.5 (Fig. 4f. Extended Data Fig. 10f). Although the power of decay did not depend on the modification, the contact frequency was consistently around two to four-fold lower for the acetylated array at the same nucleosome distances.

By calculating the insulation score [45] for the reconstructed contact map, we further confirmed the structural heterogeneity in the Center48-Kac and Every12-Kac arrays (Fig. 4g and Extended Data Fig. 10g). The score was consistently negative around the acetylated regions, indicating that the acetylated regions act as insulating boundaries.

## Discussion and conclusion

We constructed long chromatin arrays with designed histone modification patterns, demonstrating that these patterns alone can drive macroscopic changes in chromatin architecture. Isolating the impact of histone modifications without the confounding effects of other *in vivo* factors, we compared arrays of the same length and showed how acetylation induces larger, slower fluctuations. The acetylation-induced spreading, almost undetectable at the 12-mer scale, became prominent at larger sizes, indicating a length-dependent influence of acetylation on chromatin architecture.

The observed relationship between the spatial extent and the time scales of fluctuation is consistent with the Zimm model of polymer dynamics [33, 34]. Interestingly, our *in vitro* Hi-C experiments revealed a scaling exponent of − 1.5 for contact frequency decay [34], suggesting a Gaussian chain conformation. This apparent Gaussian behavior, as well as the orientation-independent contacts observed for 12-mers, is consistent with the previous observations that polynucleosomes with long linker length (nucleosome repeat length = 197 bp) without linker histone protein shows irregular structure [42]. This structure is distinct from the regular helical 30 nm fibers reported in reconstituted chromatin with linker histone H1 or shorter linker length supplied with magnesium ion [41, 42, 18, 43]. The observed −1.5 exponent is more typical of yeast chromatin [46] than of the compact organization of mammalian interphase chromatin [47], implying that our *in vitro* system mimics aspects of a more open chromatin state, even for non-acetylated arrays at physiological salt conditions.

Furthermore, the observed Gaussian behavior, implying interaction-free nucleosomes, seems to contradict our modification-dependent contact maps. This apparent discrepancy highlights the limitations of simplistic polymer models that neglect specific internucleosomal interactions. We hypothesize that transient, weak interactions, modulated by histone modifications, could be sufficient to influence contact frequencies without drastically altering the overall polymer scaling behavior observable at the resolution of our experiments. This is also consistent with the observation that the end-to-end fluctuations were indistinguishable for the two heterogeneous 96-mers with the same acetylation density, while their structures observed in the contact map were clearly distinct. Such interactions might not be strong enough to drive substantial deviations from Gaussianity but could nonetheless bias the probability of specific contacts within the chromatin fiber.

The “chromatin-only” approach establishes a baseline for understanding the contribution of histone modifications to the higher-order structure, which has previously been modeled with polymer physics with different levels of detail [48, 49, 50, 51]. Future investigations incorporating other binding components and modifications, such as H1, bromodomain-containing proteins with the tail acetylation, HP1 with H3K9 methylation, and exploring the influence of DNA-binding factors will be essential to further elucidate the complex relationship between epigenetic marks, chromatin organization, and ultimately, genome function. This controlled in vitro system allows us to systematically address the extent to which specific modifications and interacting components contribute to the diverse structural and functional landscapes of chromatin.

## Acknowledgments

We thank Y. Sakaide for technical assistance, O. Nishimura, T. Kondo (Laboratory for Developmental Genome System, RIKEN Center for Biosystems Dynamics Research) for assistance in sequencing, T. Yamamoto for the discussion on the polymer relaxation time scale, and K. Adachi, S. Kohyama, R.T. Cerbus, and I. Hiratani for reading and commenting on the manuscript. We also thank all the past and current members of the Kawaguchi lab for helpful discussions.

## Author contributions

Conceptualization: Y.T.F, K.K.; Methodology: Y.T.F, T.K., S.K., L.Y., S.Z., C.T., M.K., H.K., K.K.; Funding acquisition: H.K, T.U., K.K.; Investigation: Y.T.F, S.K., L.Y., S.Z., C.T., M.K.; Project administration: Y.T.F., K.K.; Resources: M.W., T.U.; Supervision: K.K.; Writing — original draft: Y.T.F, K.K.; Writing — checking protocols: Y.T.F, T.K., M.W., S.K., L.Y., S.Z., C.T., M.K., H.K., T.U., K.K.; Writing — review and editing: Y.T.F, T.K., H.K., T.U., K.K.

## Data and materials availability

The data and code needed to reproduce our results will be uploaded to an online repository.

**Extended Data Figure 1:**
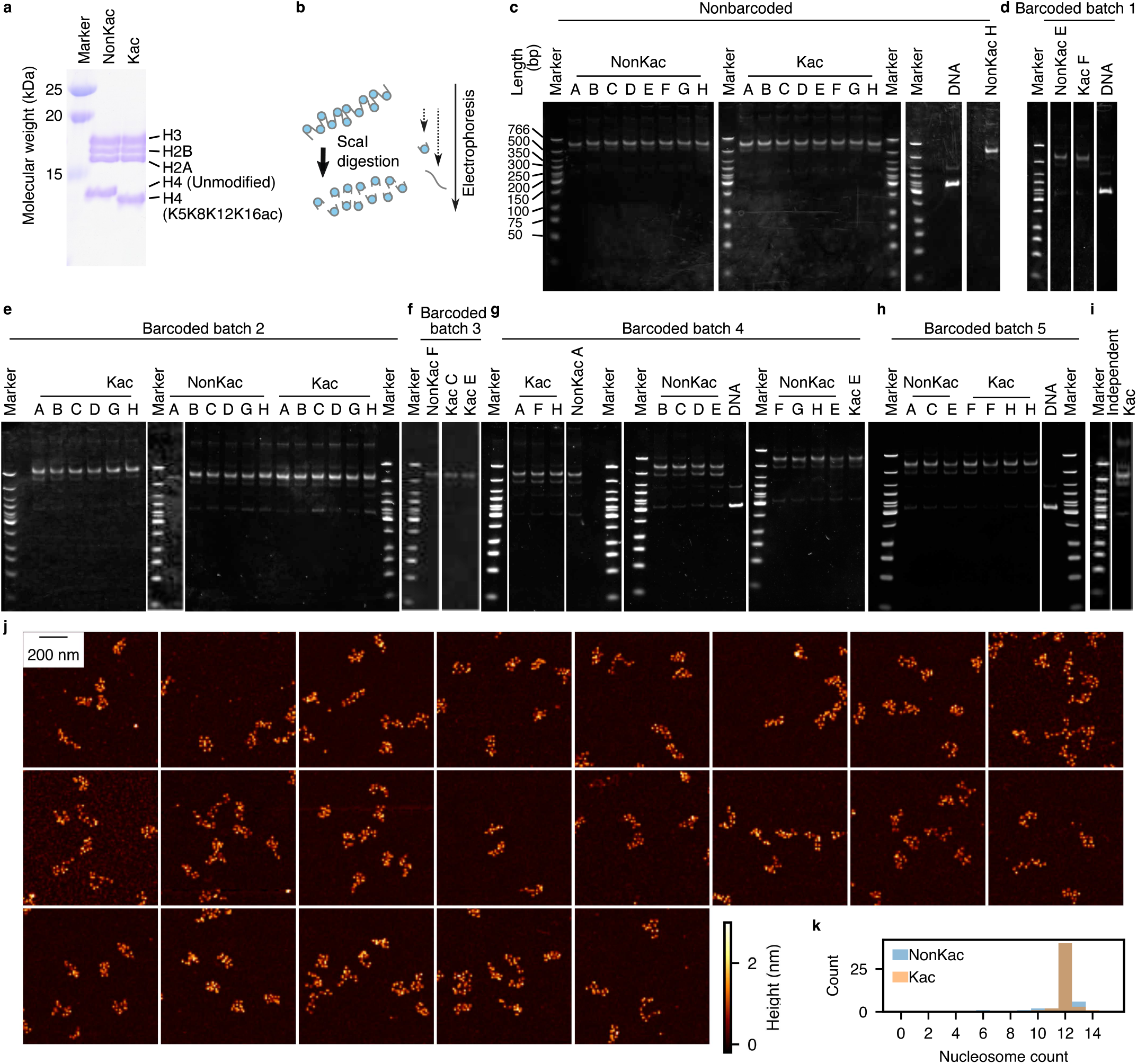
Validation of acetylated and non-acetylated 12-mer arrays. **a**. SDS-PAGE of reconstituted histone octamers with and without H4 Lysine tail acetylation. **b**. Schematic of nucleosome reconstitution confirmation by gel-shift assay after ScaI digestion. **c-i**. Confirmation of the nucleosome reconstitution by gel-shift assay for the non-barcoded (c) and barcoded (d-i) 12-mer arrays. **j**. Example snapshots of AFM with reconstituted 12-mer arrays with barcode. **k**. Distribution of the nucleosome counts in the clusters found in the AFM images.

**Extended Data Figure 2:**
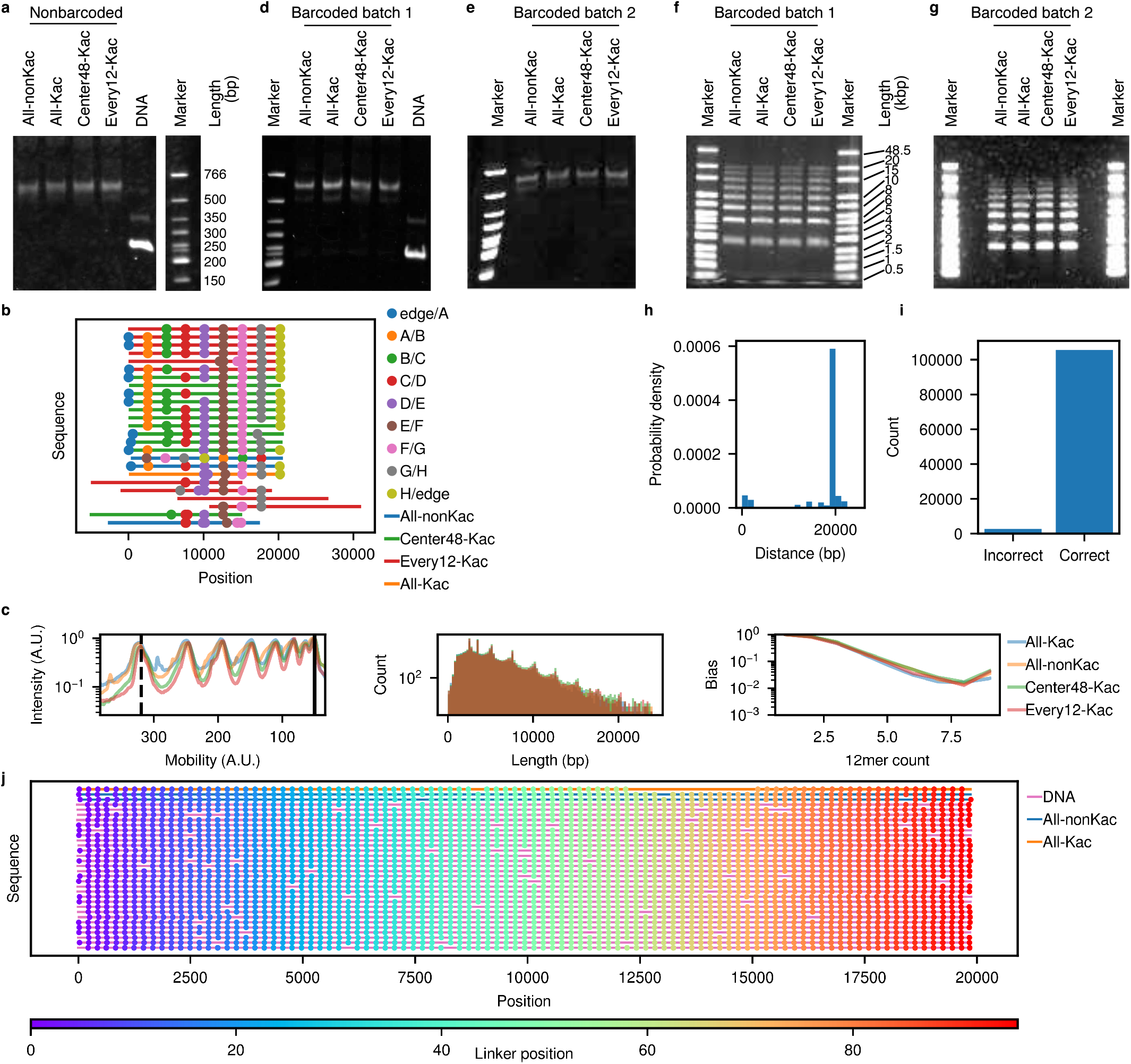
Validation of ligated arrays. **a**. Confirmation of the nucleosome reconstitution by gel-shift assay for the non-barcoded ligated arrays. **b**. Each sequence for Fig. 1e. **c**. Illustration of nanopore sequencing bias. The left, center, and right plots show the fluorescent intensity in 0.3% agarose gel electrophoresis, the length distribution of the sequences, and the estimated read bias, respectively. In the left plot, the black dashed and solid lines show the mobility for the 12-mer and 96-mer, respectively. The plots show the data for experiment 2 for the non-barcoded array (Table S1). **d**,**e**. Confirmation of the nucleosome reconstitution by gel-shift assay for the barcoded ligated arrays. **f**,**g**. Electrophoresis result of the ligated product of eight barcoded 12-mer arrays. **h**. Sequence bias-corrected distribution of the length of barcoded DNA when subsampling the sequences that include both ends of the 96-mer design. **i**. The correct and incorrect ligation count at the junctions for the barcoded arrays. **j**. Aligned sequences of the barcoded arrays with a length close to 96-mer.

**Extended Data Figure 3:**
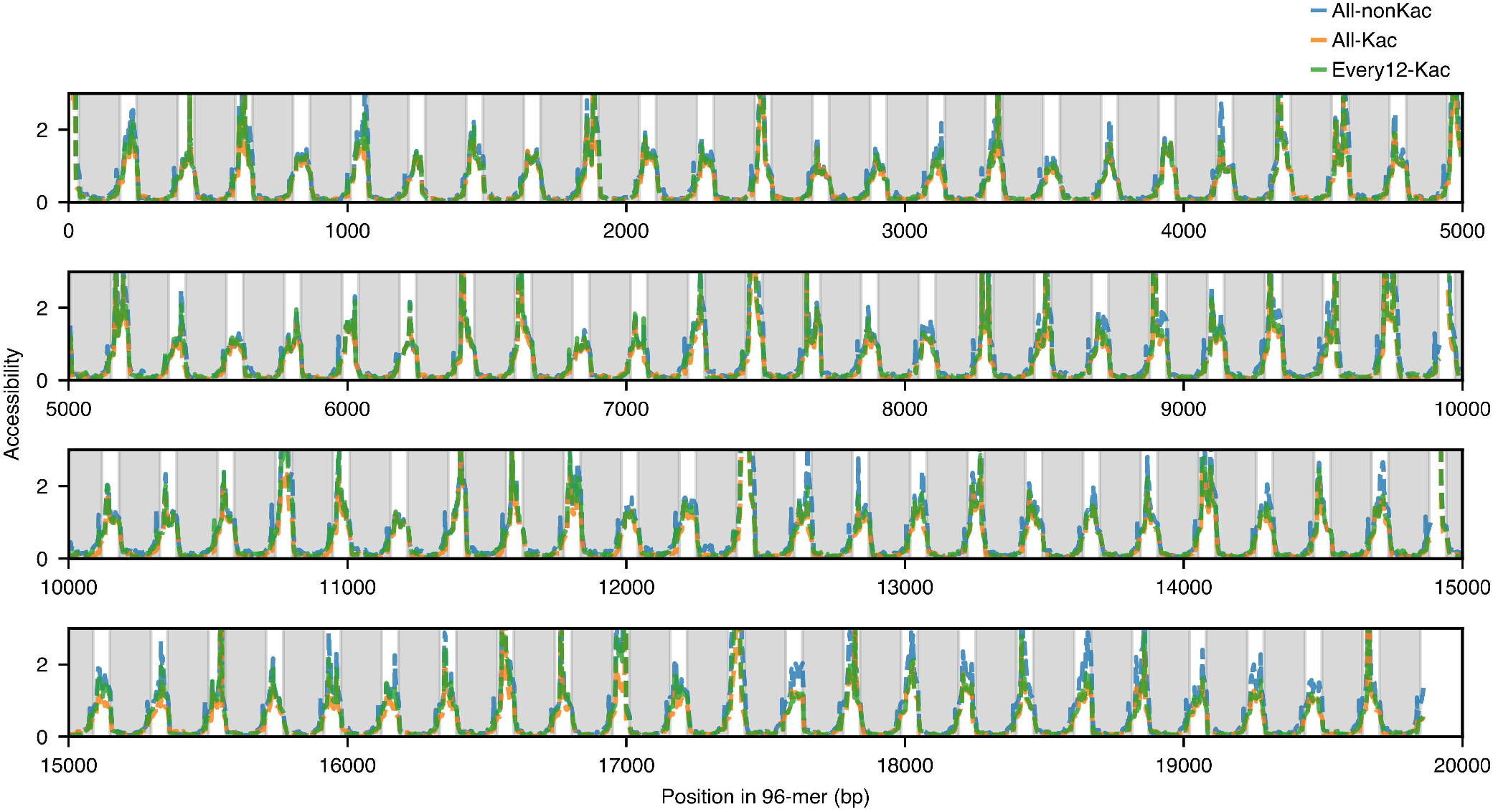
Accessibility assay of the ligated arrays. The accessibility was measured as the ratio of the fraction of adenine methylated by EcoGII for the ligated nucleosome arrays to that of DNA without histones.

**Extended Data Figure 4:**
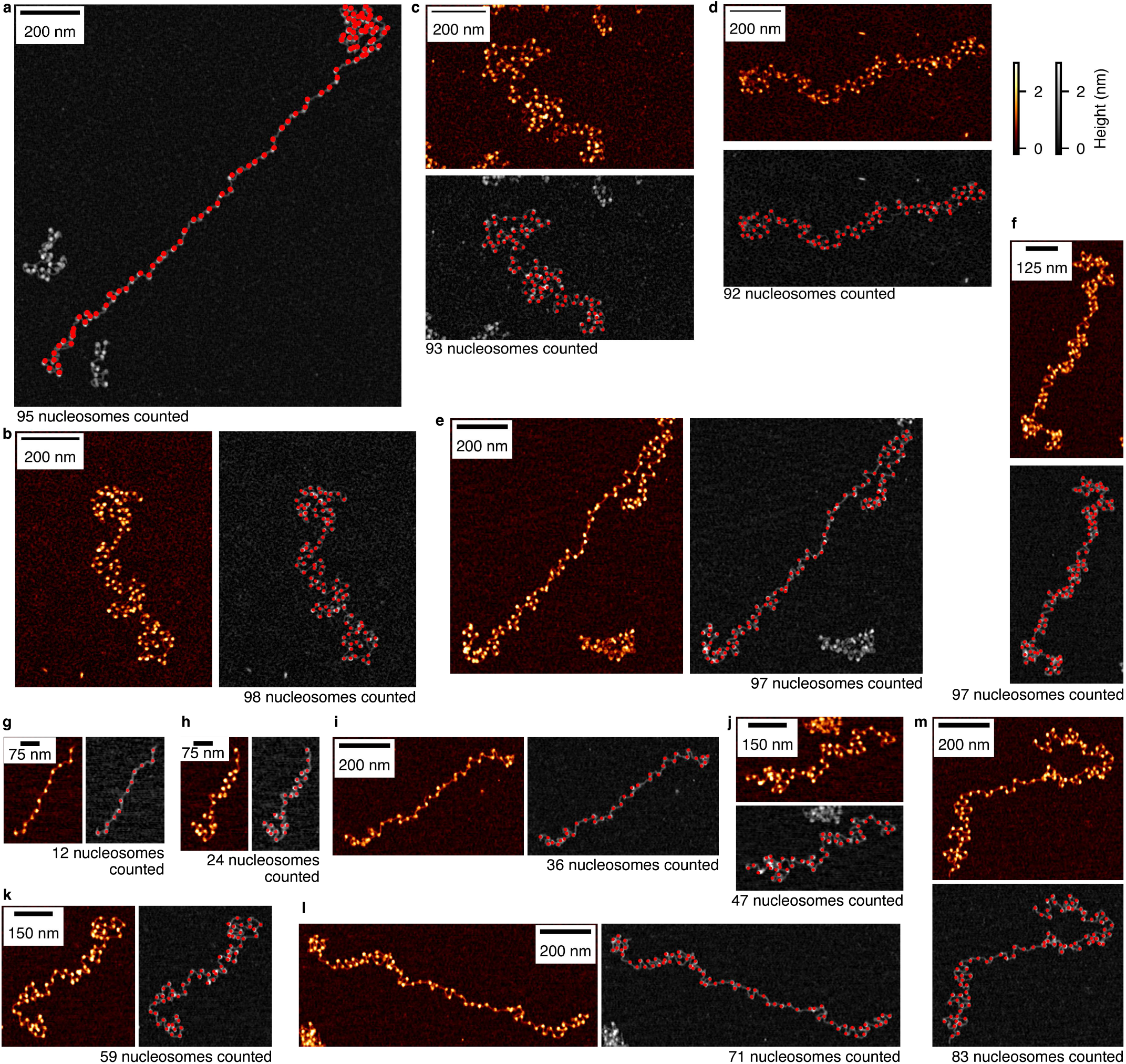
AFM images and nucleosome counts of reconstituted arrays. **a-f**. Arrays with nucleosome counts close to 96. **g-m**. Arrays with nucleosome counts close to 1 × 12, …, 7 × 12. All-nonKac (a,i,k-m), All-Kac (b,f) Center48-Kac (c), and Every12-Kac (d,e,g,h,j) samples. Barcoded (a,e-m) and non-barcoded (b-d). The raw image for (a) is shown as Fig. 1i. The red points show the detected nucleosome positions.

**Extended Data Figure 5:**
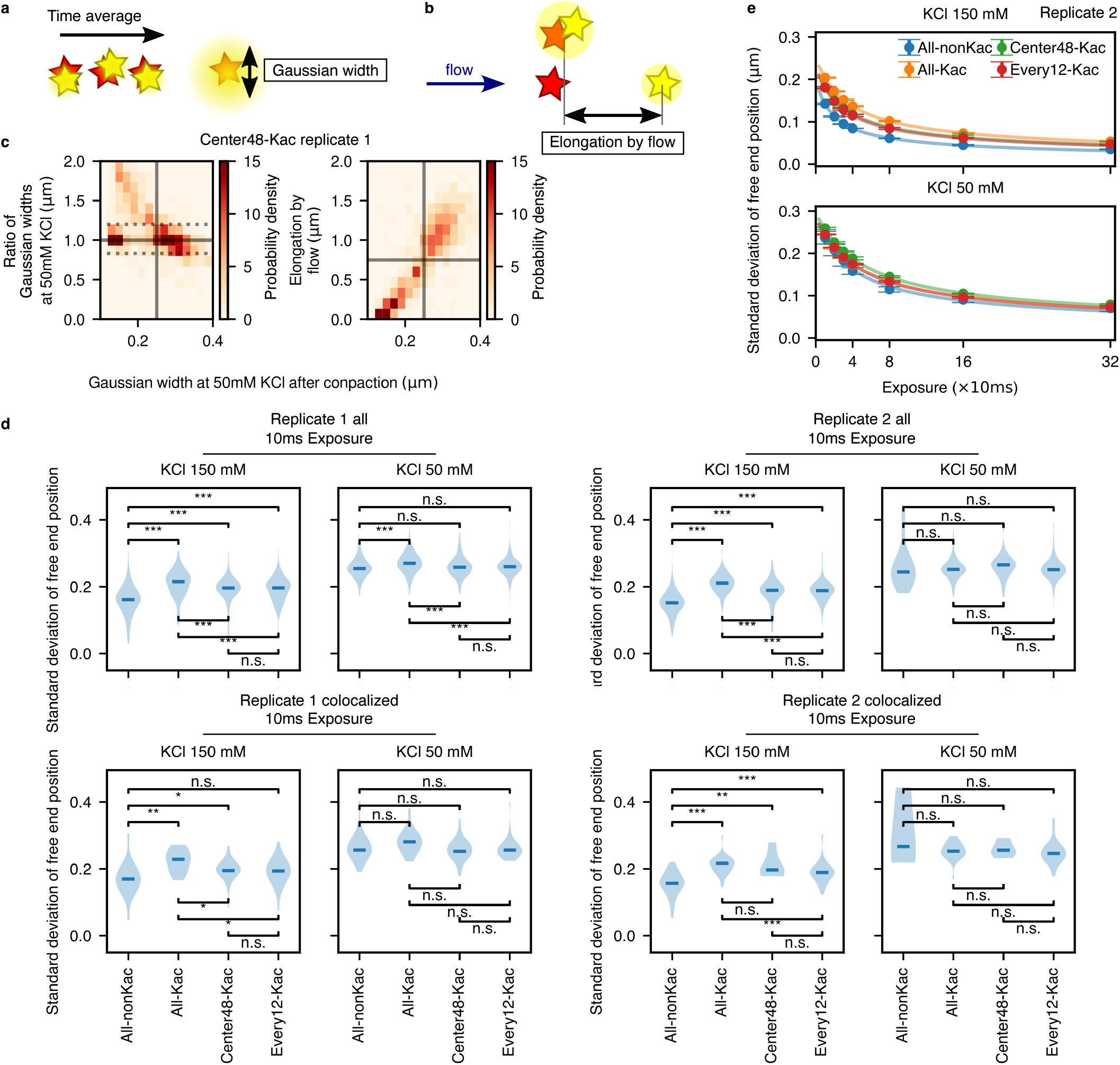
Filtering scheme and replicate results for single-molecule microscopy. **a**,**b**. Schematic for the definition of the Gaussian width (a) and the elongation by flow (b). **c**. (left) 2D histogram of the Gaussian width at 50mM KCl (after compaction by 150mM KCl) and the ratio of the Gaussian widths before and after the compaction. (right) 2D histogram of the Gaussian width at 50mM KCl after the compaction and the elongation by flow. The solid gray line shows the thresholds for the Gaussian width after compaction and elongation by flow. The gray dotted lines show the thresholds for the ratio of Gaussian widths before and after the compaction. The plots show data for the Center48-Kac sample (replicate 1). **d**. Replicate results for Fig. 2d, plotted using all the filtered free end tracks (top) and only tracks that colocalize with the tethered end fluorescent signals (bottom). The asterisks indicate the statistical significance of the Mann-Whitney test adjusted by the Benjaminini-Yekutieli method: “*”, “**”, and “***” correspond to statistical significance with *p* < 0.05, *p* < 0.01, and *p* < 0.001, respectively. “n.s.” means not significant. **e**. Replicate results for Fig. 2e.

**Extended Data Figure 6:**
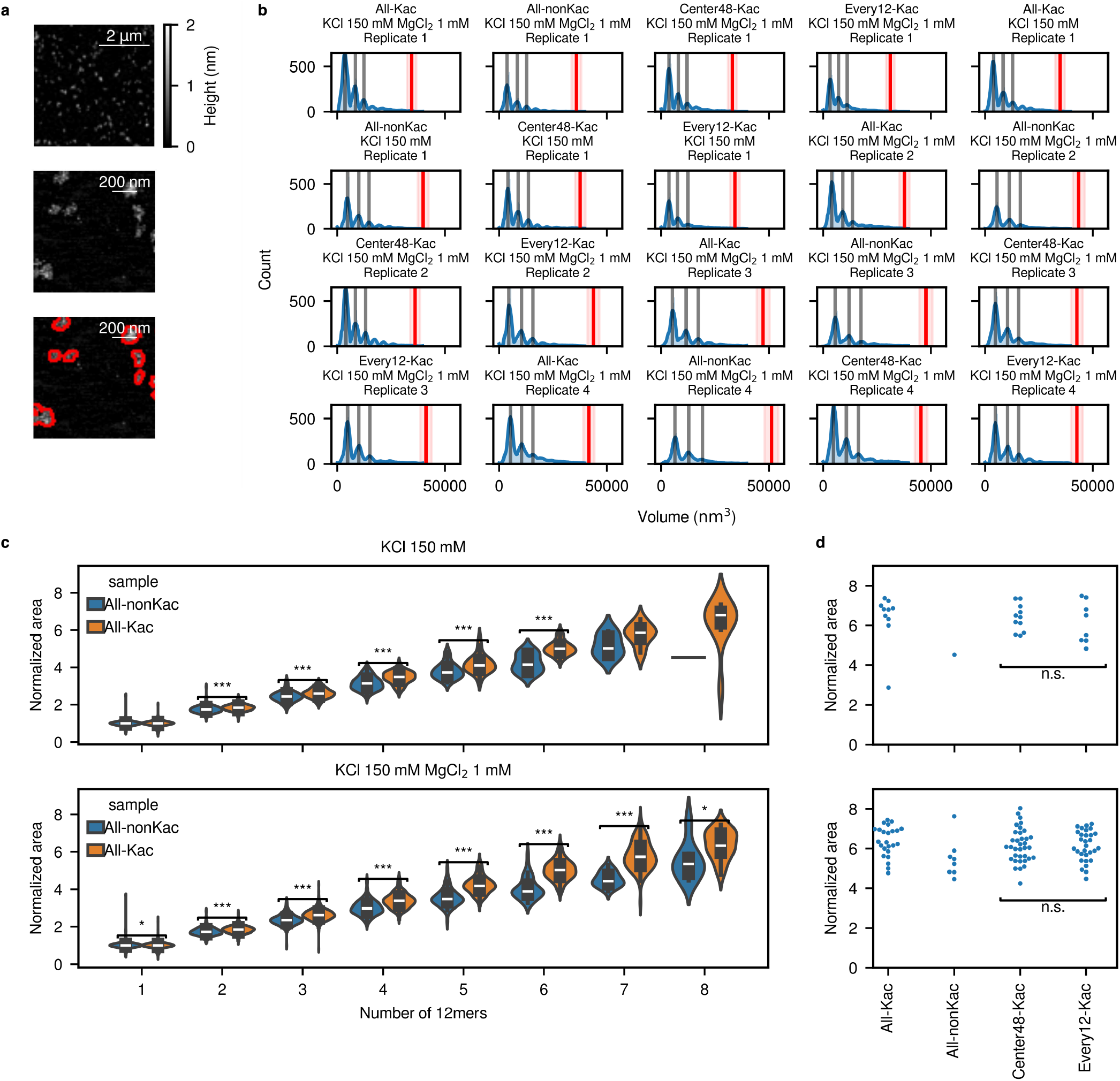
Size measurement of ligated products using AFM. **a**. Representative low-resolution AFM images of ligated products. Red lines indicate the boundaries of the detected blobs. **b**. Histograms showing the measured volumes of the blobs. Gray lines correspond to the estimated volumes for 12-mer, 24-mer, and 36-mer. The red line and shaded region show the estimated volume and volume range of the 96-mer. **c**. Blob area normalized to that of the 12-mer, grouped by the count of the 12-mers estimated by the volume. **d**. Pattern-dependent normalized areas for arrays whose volumes are in the estimated range for the 96-mer.

**Extended Data Figure 7:**
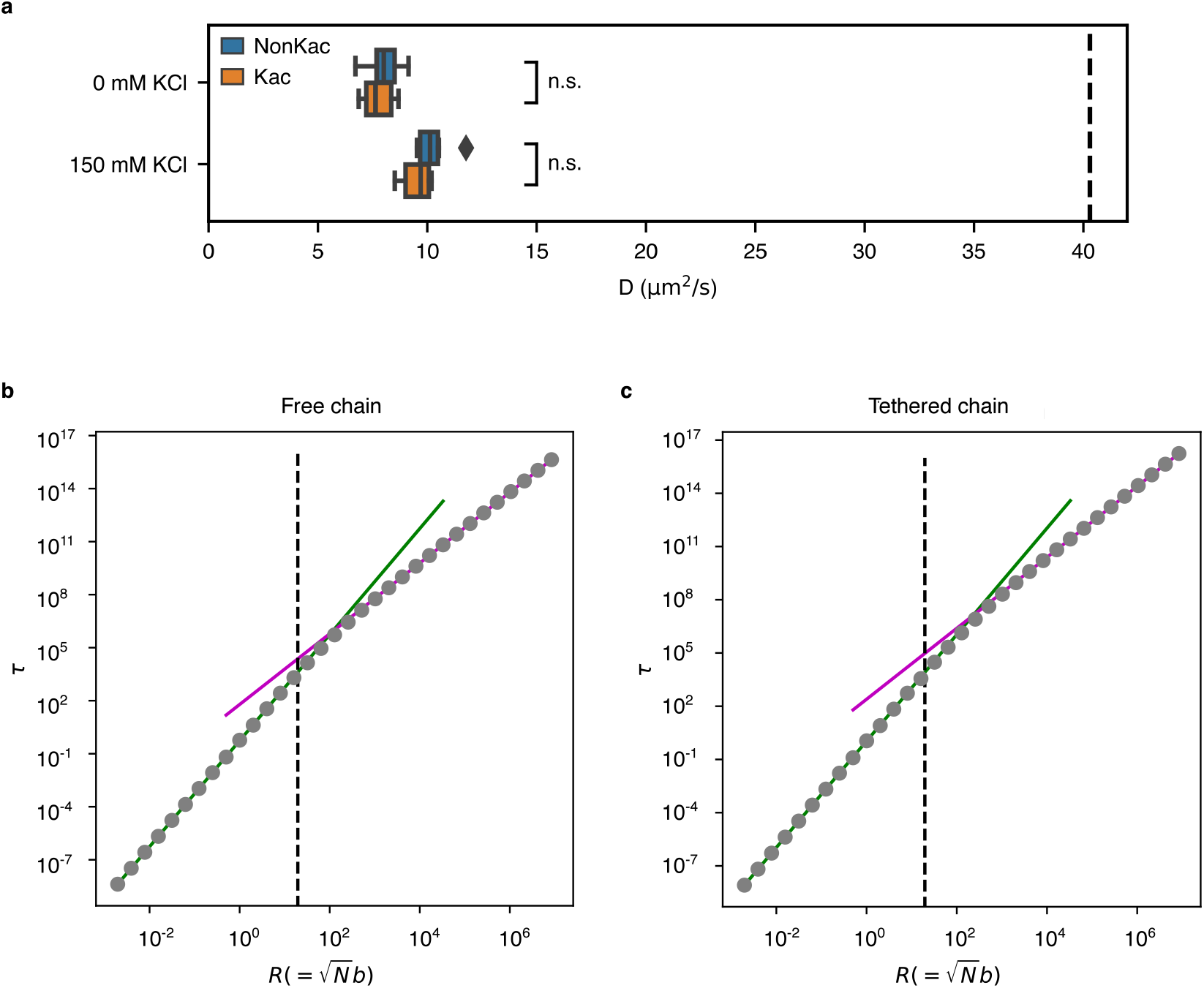
Comparison between experimental data and Zimm model. **a**. Diffusion constant of the 12-mer arrays measured by single-molecule tracking. The black dashed line indicates the value for mononucleosome reported in [1]. The value of the 12-mer diffusion constant is around one-fourth or less compared with a mononucleosome. “n.s.” indicates a nonsignificant difference according to the Mann-Whitney test. **b**,**c**. Zimm model numerics for prefactor estimation regarding the relation between size 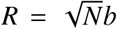 and time scale *τ* (see section ‘Zimm model with tethered end’ in the supplementary). The Kuhn length *b* is varied with fixed *a* = 1 as well as *η* = *k*_B_*T* = 1, and *N* = 96. Dashed line is at *b* = 2. Green lines are extensions of fits using the two left-most points to obtain α, and the magenta lines are fits using the two right-most points for *v*. **b**. Free chain case. Green line: *τ* = 0.5501*R*^2.9998^. Magenta line: *τ* = 1.9993*R*^2.0000^, where the exponents and the prefactor are both fitting parameters. **c**. Tethered chain case. Green line: *τ* = 1.0857*R*^2.9998^. Magenta line: *τ* = 8.0771*R*^2.0000^, where the exponents and the prefactor are both fitting parameters.

**Extended Data Figure 8:**
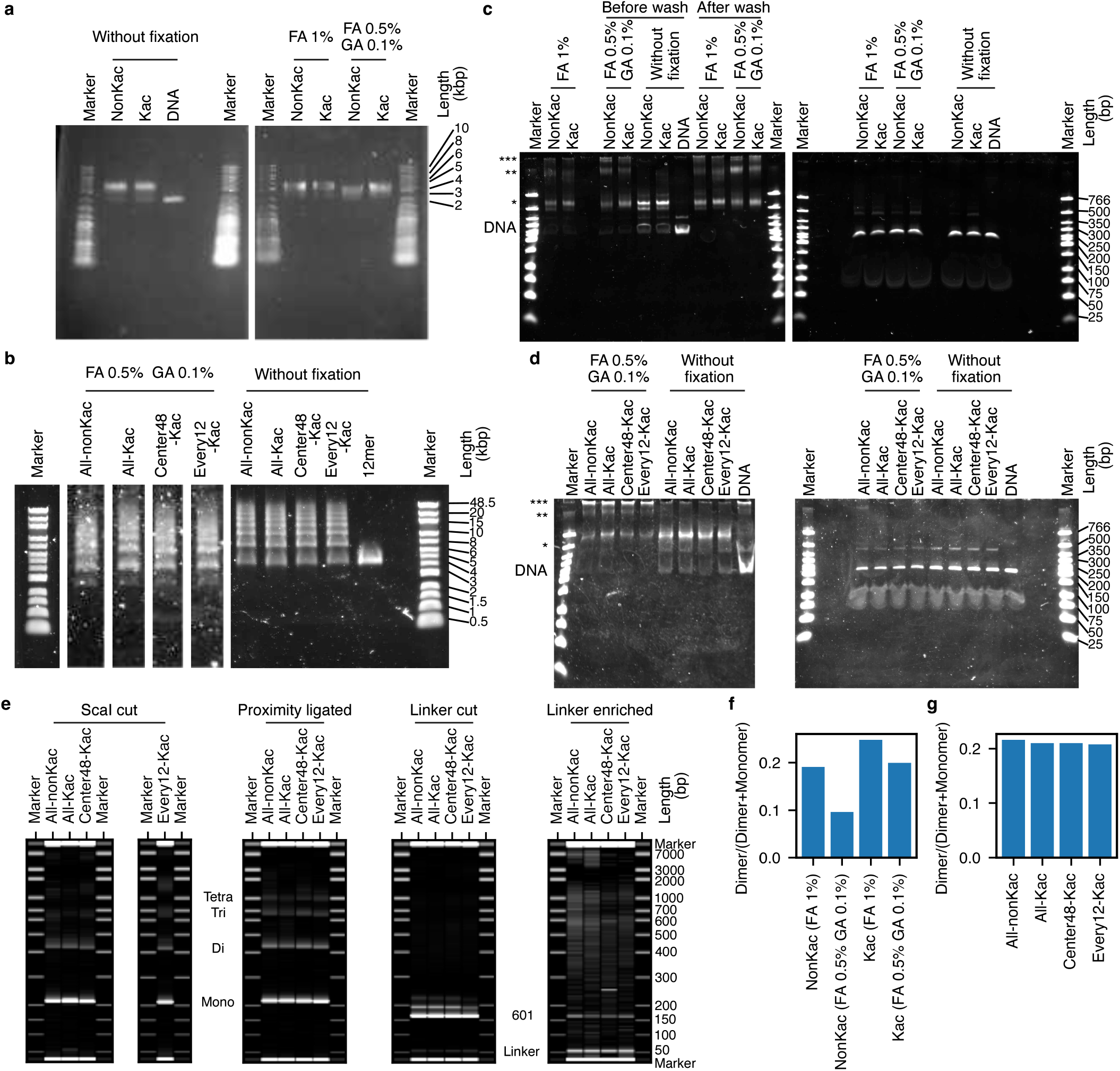
Experimental details of the contact map experiment. **a**. 0.7% agarose gel electrophoresis in 0.2X TBE of the 12-mer samples before and after fixation. **b**. 0.3% agarose gel electrophoresis in 0.2X TBE of the ligated samples before and after fixation. **c**,**d**. 5% native PAGE in 1X TBE of 12-mer (c) and ligated (d) samples after ScaI digestion. The left and right gels correspond to the samples before and after SDS/PK digestion, respectively. The band positions for the bare DNA, mononucleosome, dinucleosome, and products with lower mobility are shown by the text “DNA” and the single, double, and triple asterisks, respectively. **e**. Pseudogel images showing the Bioanalyzer traces in each protocol step (see Methods). In the left images, the band positions for the single, double, triple, and tetra nucleosomes are shown, respectively, by the text “Mono”, “Di”, “Tri”, and “Tetra”. In the right images, the band positions for the Widom 601 and linker sequences are shown by the texts “601” and “Linker”. **f**,**g**. Relative abundance of dinucleosomes after ScaI digestion, calculated from the bioanalyzer traces. 12-mer samples (f) and ligated samples (g). FA and GA abbreviate formaldehyde and glutaraldehyde.

**Extended Data Figure 9:**
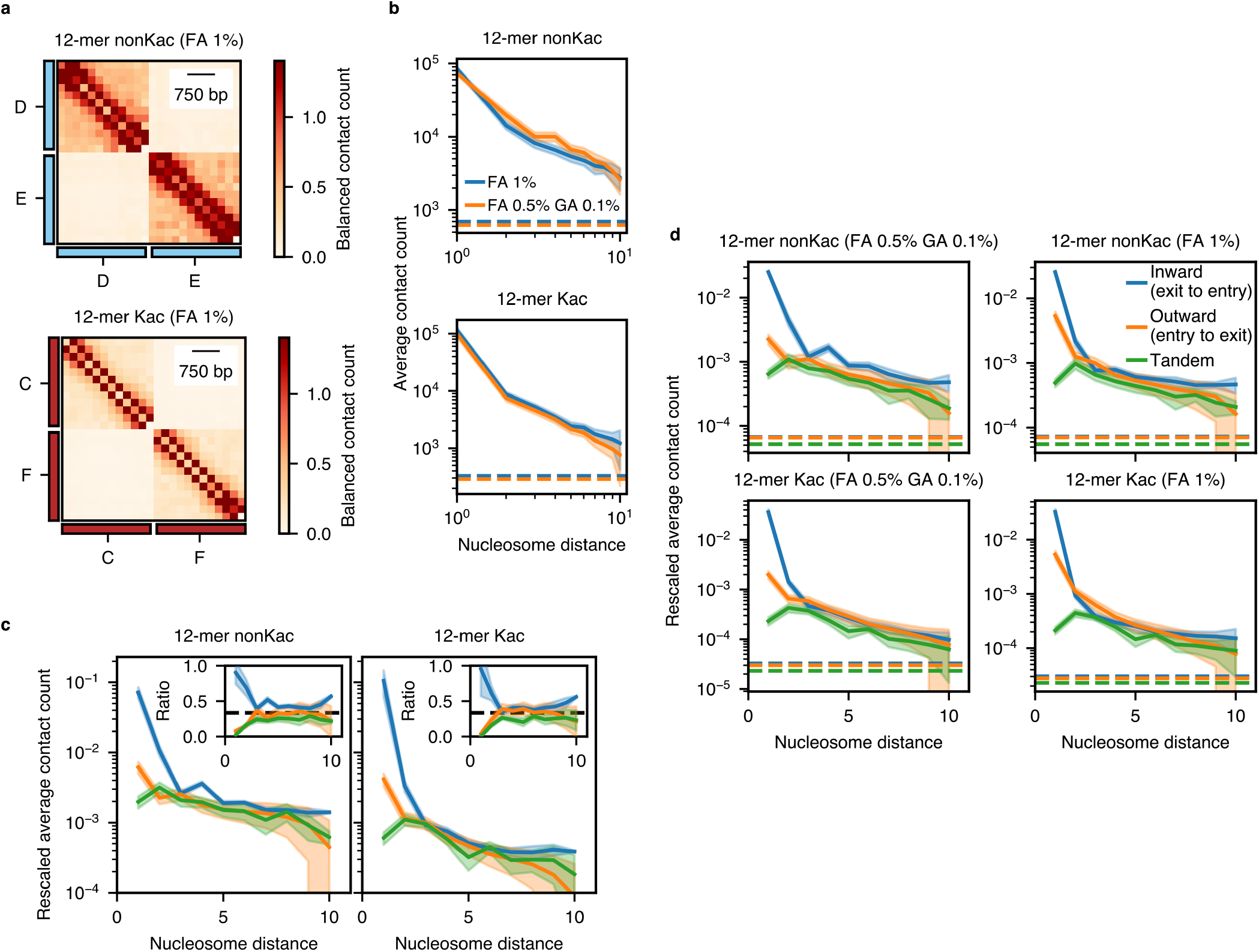
Fixation method dependence on *in vitro* Hi-C. **a**. Contact map obtained from mixed 12-mers with distinct barcodes fixed with 1% of formaldehyde (FA). The contact is balanced and normalized so that the average contact frequency is 1. **b**. Contact frequency as a function of the nucleosome position with different fixation methods. The shaded area represents the uncertainty (twice the s.e.m.). The dashed lines indicate the average contact frequency of the barcodes originating from the different 12-mer arrays. **c**. The same plot as Fig. 3f for the balanced matrices. **d**. Raw data for Fig. 3f and the same plot for the samples fixed with 1% FA. The shaded area represents the uncertainty (twice the s.e.m.). The dashed lines indicate the background level, estimated as the average contact frequency of the barcodes originating from the different 12-mer arrays.

**Extended Data Figure 10:**
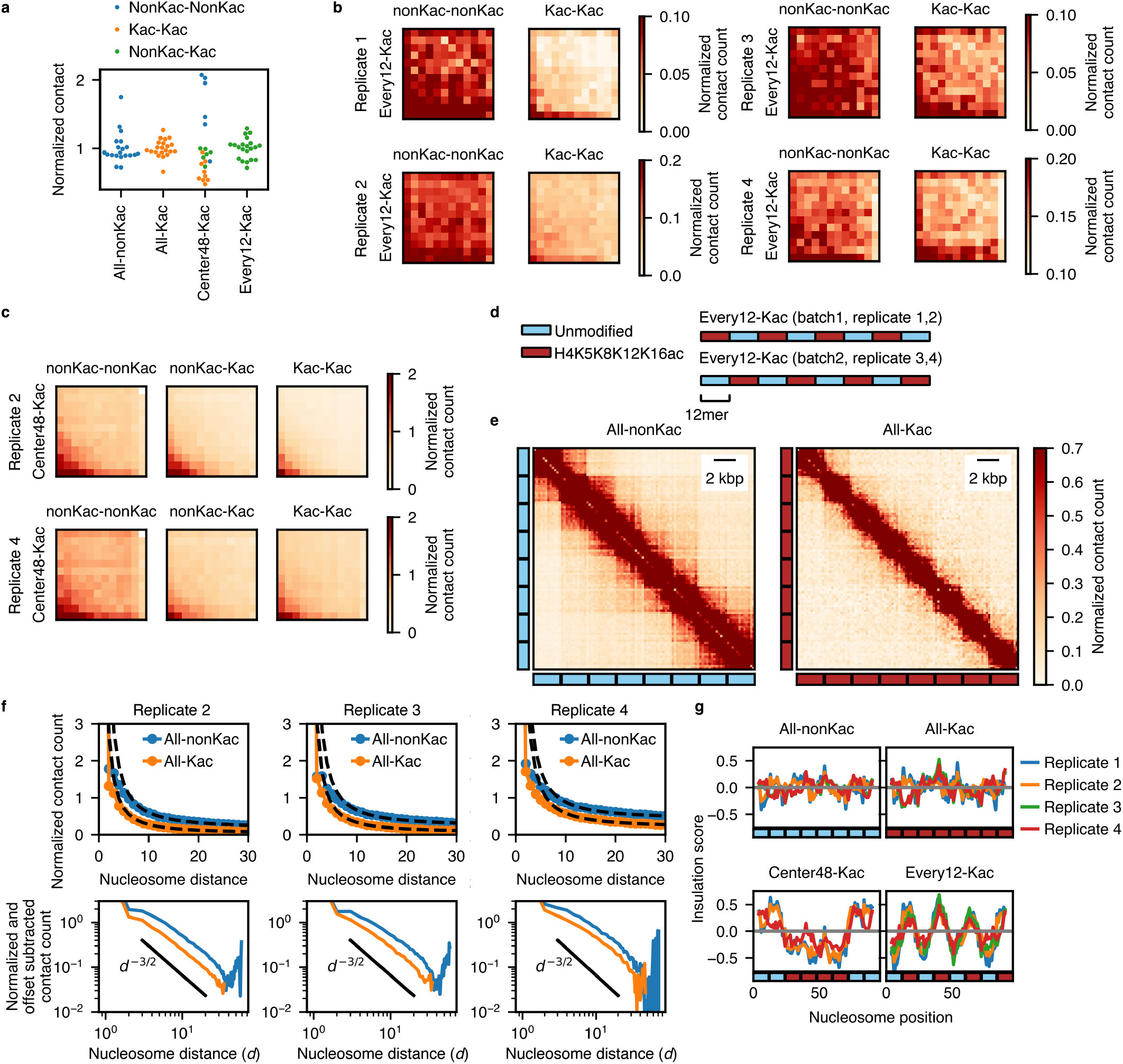
Additional results of *in vitro* Hi-C for ligated arrays. **a**. All data for Fig. 4c, categorized by the nucleosome type. **b**. Accumulated contact frequency between second adjacent 12-mer regions for the Every12-Kac arrays. **c**. Replicate results for Fig. 4d. **d**. Design of histone modification patterns for the Every12-Kac arrays in each replicate. **e**. All-to-all contact map of the ligated nucleosome arrays with homogeneous patterns. **f**. Replicate results for Fig. 4f. **g**. Replicate results for Fig. 4g with the data for homogeneous arrays. The sequence of replicates 2 and 3 for Every-12 arrays are flipped to match the modification patterns (See (d)).

## Methods

### DNA design and preparation

All LB plates and medium were supplemented with 0.1 mg/mL ampicillin. Buffers recommended by New England Biolabs were used for the restriction enzyme digestion and ligation reactions, if not specified.

For the non-barcoded arrays, DNA sequences contain 12 repeats of the Widom 601 nucleosome positioning sequence [2] separated by a 61 bp linker DNA. The two ends of the whole 12-mer sequence were designed to be cut by the BglI restriction enzyme, so that the 3 bp sticky ends differ from each other by at least 2 bp, except for the 5’ end of A and the 3’ end of H, which ligate to the short labeled oligonucleotides (Fig. 1b). The transformed *E. coli* cells (New England Biolabs, C2987) were plated on LB agar plates and incubated overnight at 37 °C. Colonies were picked and grown in 3mL LB medium for 7-9 hours. The cells were then transferred to 2.5 L LB medium and cultured overnight at 37 °C. Plasmids were extracted using the Qiagen Plasmid Giga Kit (Qiagen, 12191) and digested with 0.1 − 0.4U/µg BglI (New England Biolabs, R0143) to excise the insert from the backbone, and 0.15 − 0.2U/µg EcoRV-HF (New England Biolabs, R3195) to digest the backbone into 200-300 bp fragments for two to three overnights. The 12 × 601 fragments were then purified by two rounds of polyethylene glycol (PEG) precipitation (4.33% polyethylene glycol 6000 (Nacalai Tesque, Inc., 28254-85) and 0.595M NaCl (Nacalai Tesque, Inc., 31320-05) final) followed by ethanol precipitation. The DNA was then resuspended in TE buffer (Nacalai Tesque Inc., 32739-31) and stored at − 20 °C.

The plasmids with the barcoded arrays were constructed to have 60 bp linker DNA, including two 17 bp barcodes, with a digestion site by ScaI in between (Fig. 3a). The 12 × 601 fragments were prepared similarly to the non-barcoded plasmids.

### End labeling of A and H fragments

Following a method in Ref. [3], modified oligonucleotides 5’-Ph-TAGTCTGC(T*)CAGATATCGTCG-Biotin-3’ and 5’-CACAGCTG(T*)CTGAG-3’, where T* represents the amine-linked dT with the C6 linker (purchased from Merck), was labeled by Atto647N-NHS ester (ATTO-TEC GmbH, AD647N-31) and Atto565-NHS ester (ATTO-TEC GmbH, AD 565-31), respectively. Those fragments were annealed to the complementary oligonucleotides and ligated to the “A” and “H” arrays, respectively. See Supplementary Methods for details.

### Protein expression and purification

Recombinant histone H4 with or without residue-specific acetylation at K5/K8/K12/K16 was synthesized by genetic code reprogramming in a coupled transcription-translation cell-free system. The cDNA containing human histone H4 with or without the codons of K5/K8/K12/K16 replaced with the TAG triplets and a terminal TAA stop codon were subcloned into a pCR2.1-TOPO plasmid (Thermo Fisher Scientific, K450002). The unmodified and K5/K8/K12/K16-tetra-acetylated histone H4 (H4K5K8K12K16ac) proteins were designed to have an N-terminal histidine-rich affinity tag (N11; MKDHLIHNHHKHEHAHA) followed by a TEV protease recognition sequence (EHLYFQ), which has no additional N-terminal sequence after cleavage by TEV protease. Protein synthesis in the cell-free system was performed as described [4], using the subcloned plasmids, with a 9-mL reaction solution dialyzed against a 90-mL external feeding solution at 37 °C for 6–18 hours. Precipitates of the unmodified H4 and H4K5K8K12K16ac were separated by centrifugation for 30 min at 30, 000 × *g* at 4 °C.

Unmodified human histones were recombinantly expressed in *E. coli*. The cDNA containing human histone H2A type 1-B/E, H2B type 1-J, and H3.1 were subcloned into a pET15b-derived plasmid containing an N-terminal 6×His tag (MGSSHHHHHHSSG) in which the thrombin cleavage site of pET15b (Merck Millipore, 69661) was replaced by the TEV protease recognition sequence. The H2B expression construct contains a glycine residue at the end of the TEV protease recognition sequence which remains after cleavage by TEV protease. The H2A and H3 expression constructs have no additional N-terminal sequence after cleavage by TEV protease. These histones were expressed in LB broth of *E. coli* BL21 (DE3) cells at 37 °C. Induction of the recombinant histone proteins, separation of their precipitates, and preparation of the unmodified and H4K5K8K12K16ac-containing histone octamers *in vitro* were performed essentially as described [4].

### Reconstitution of 12-mer nucleosome arrays

The template DNA hosting 12 repeats of the 601 sequence was mixed with the reconstituted histone octamer and short octamer sink DNA fragments with the molar ratio to the 601 sequences of 1.2 and 5/12 for the non-barcoded arrays and 1.1 and 1/3 for the barcoded arrays, respectively. The reconstitution was performed by the salt-gradient dialysis from 2M KCl to 250mM KCl and then to No-salt buffer (10mM Tris-HCl (pH 7.5) and 1mM EDTA) followed by purification by 10% - 40% sucrose gradient ultracentrifugation. Reconstituted chromatin was routinely checked by the band shift assay in agarose gel electrophoresis and PAGE after ScaI digestion in the ScaI cut buffer (10mM Tris-HCl (pH 7.5), 0.5mM MgCl_2_ and 50mM NaCl). See Supplementary Methods for details.

### Ligation of 12-mer arrays into 96-mer nucleosome arrays

The frozen stocks of the 12-mer arrays were quickly thawed by hand warming and then placed on ice. The equal amount of 12-mer arrays (1-2.5 µg for each) were mixed and purified again by 10% - 40% sucrose gradient. The arrays were then further concentrated to 0.4-1.1 µg/µL by centrifugation in Vivacon 500 100kDa (Sartorius, VN01H42). The 12-mer mixture was ligated in the final concentration of 0.3-0.52 µg/µL (fixed to the same concentration for the same batch) with 133U/µL T4 ligase (New England Biolabs, M0202M) and 50mM Tris-HCl (pH 7.6), 2mM ATP (New England Biolabs, P0756S), 3mM MgCl_2_, 10mM DTT and 0.01mg/mL BSA (New England Biolabs, B9200S) at 16 °C overnight.

The ligated products were then washed three times with TEN 50 buffer (10mM Tris-HCl (pH 7.5), 1mM EDTA and 50mM NaCl) in the 100 kDa MWCO Amicon Ultra centrifugal filter at 14, 000 × *g* for 3min (15 min for the last wash) and once with the final buffer (No-salt buffer for the non-barcoded array and the TE0.1D buffer (10mM Tris-HCl (pH 7.5), 0.1mM EDTA and 1mM DTT) for the barcoded array) for 15min. The final ligated product was stored at 4 °C for a maximum of six weeks (non-barcoded arrays) and for seven weeks (barcoded arrays) with little nucleosome dissociation (verified by ScaI digestion and native PAGE as described).

The ligated arrays were checked by native PAGE after ScaI digestion as described (Extended Data Fig. 2a,d,e). The ligation was confirmed by digesting the ligated product by SDS/PK (0.5 mg/mL Proteinase K (FUJIFILM Wako Pure Chemical Corp., 169-21041 or 166-28913) 10mM Tris-HCl (pH 8.0), 40mM EDTA and 0.125% SDS) and performing electrophoresis with 0.3% agarose in 1X TAE (Fig. 1d and Extended Data Fig. 2f,g).

### Validation of the ligation order by nanopore sequencing

SDS/PK-digested samples were purified using Ampure XP beads (Beckman Coulter, A63880) and sequenced using the Flongle or MinION flow cell (Oxford Nanopore Technologies). The input amount, purification condition, sequencing kit, and flow cell are summarized in Table S1.

### Accessibility assay for reconstituted chromatin

Chromatin samples and equimolar mixture of the bare DNA fragments were respectively mixed with 50mM Tris-HCl (pH 7.5), 150mM KCl, 1mM MgCl_2_, 0.1mg/mL BSA (New England Biolabs, B9000S), 2.4mM S-adenosylmethionine (New England Biolabs, B9003S) and 0.645U EcoGII methyltransferase (New England Biolabs, M0603S) at the concentration of 4 ng/µL in the final volume of 33 µL, and incubated at 37 °C. The reaction was quenched by taking 10 µL of the sample after 10 min and 40 min and adding 2.5 µL of 6 × SDS/PK buffer (60mM Tris-HCl (pH 8.0), 240mM EDTA, and 3% SDS), 1.5 µL of 5mg/mL Proteinase K, and 1 µL of ultrapure water and incubating at 65 °C (lid 75 °C) for more than 2 hours. 10 µL of the samples were immediately quenched after mixing (“0 min” sample). DNA was purified by two rounds of purification by the Serapure magnetic beads (×1.8 volume to the sample; See Supplementary Methods), processed using the library preparation kit SQK-NBD114.24 (Oxford Nanopore Technologies) and sequenced using the MinION R10.4 flowcell (Oxford Nanopore Technologies, FLO-MIN114).

### Observation by atomic force microscopy

For the nucleosome counting (Fig. 1 and Extended Data Figs. 1, 4), samples were diluted to 0.25-1 ng/µL and fixed with 0.1% glutaraldehyde (Nacalai Tesque Inc., 17003-92) or 0.1% formaldehyde (FUJIFILM Wako Pure Chemical Corp., 064-00406) and 0.05%glutaraldehyde on ice for 10-30 min. For the size measurement (Extended Data Fig. 6), ≈ 20 ng of the 96-mer samples (barcoded, batch 1) were diluted in the H30D buffer and the equivalent volume of H30K150 (salt2X) buffer (30mM HEPES-KOH (pH 7.5), 300mM KCl and 1mM DTT) or H30K150M1 (salt2X) buffer (30mM HEPES-KOH (pH 7.5), 300mM KCl, 2mM MgCl_2_ and 1mM DTT) were added and mixed by tapping.

The samples were mounted on a mica plate coated by poly-L-ornithine solution (Sigma-Aldrich, P3655) and observed with the Innova atomic force microscope (Bruker) using the SNL10-C probe in the tapping mode with the pixel size of 9.77 nm (size measurement) or 1-2 nm (nucleosome counting). For Fig. 1h,i and Extended Data Figs. 4a,e-m, chromatin was elongated by flow in the mounting step. See Supplementary Methods for details.

### Single-molecule observation by fluorescence microscopy

For the chamber construction for single-molecule imaging, we followed a protocol in Ref. [3] with modifications. Imaging buffer (IB) was prepared according to Ref. [3, 5]. See Supplementary Methods for details.

The chambers were washed sequentially with 200 µL of NaB buffer (pH 8.5) and T50 (pH 8.0) buffer (10mM Tris-HCl (pH 8.0) and 50mM NaCl), and incubated with 80 µL of 0.25mg/mL Neutravidin (Thermofisher Scientific, 31000) in T50 (pH 8.0) for 15min. The chambers were then washed again with 200 µL of T50 (pH 8.0), 1mL of T150 (10mM Tris-HCl (pH 7.5) and 150mM NaCl), 200 µL of T50 (pH 7.5, 10mM Tris-HCl (pH 7.5) and 50mM NaCl), and finally with 100 µL of IB (KCl 50mM). Chromatin samples, diluted to 100 pM in IB (KCl 50 mM), were injected into the chambers and incubated for 2-5min. The chambers were then washed twice with 50 µL of IB (KCl 50 mM) and observed sequentially in (1) IB (KCl 50 mM) without flow (2) IB (KCl 150 mM) without flow (3) IB (KCl 50 mM) without flow and (4) IB (KCl 50 mM) with flow (200 µL/min). After sample injection, buffer exchange was performed at the rate of 25 µL/min. 100 µL of each buffer was injected into the reservoir, from which 50 µL was introduced into the chamber by syringe pump aspiration.

The sample was observed with a microscope equipped with the objective lens UPlanApo 60x / 1.50 Oil HR TIRF (Olympus), epi-illumination by the laser light from OBIS 561nm LS (Coherent) and OBIS 637nm LX (Coherent), and the Multisplit V2 (Teledyne Photonics) for simultaneous dual-color observation by the Highly Inclined and Laminated Optical Sheet (HILO) microscopy [6]. The laser intensity was 10-20mW (561nm) and 10mW (640nm), respectively. Fluorescence images were captured using a Prime BSI sCMOS camera (Teledyne Photonics) in the Correlated Multi-Sampling mode and 10 ms exposure. The pixel size was 0.109 nm. We took movies of 100 frames in 25 fields of view with × 53.76 µm 53.76 µm area for each condition, yielding 1000-4000 fluorescent spots in total. The microscope, syringe pumps, and valves were controlled by Micro-Manager [7] and Pycro-Manager [8]. See Supplementary Methods for details.

### Diffusion coefficient measurement of 12-mer by single-molecule tracking

The Atto-565-labeled 12-mer array (barcoded “H”) was diluted to 0.2 nM in H30D buffer (30mM HEPES-KOH (pH 7.5, Nacalai Tesque Inc., 02443-34 and 28616-45) and 1mM DTT) and mixed with an equal volume of buffer with a 2× target salt concentration (H30K150 (salt2X) buffer or H30D buffer) in a well of a 96-well glass-bottom plate (Matsunami Glass, GP96003). The total volume was 50 µL. The sample was equilibrated to 30 °C in a stage-top incubator for 15min and observed with the HILO microscope as previously described. The fluorescent molecules were tracked with TrackMate [9, 10], and the blob positions were refined as the weighted centroids of background-subtracted pixel intensities in a circle with a diameter of 10 pixels around the detected point. The diffusion coefficients were then measured using the covariance-based estimator described in [11].

### *In vitro* Hi-C experiment

The chromatin conformation capture experiment was performed similarly to Ref. [12] with major differences in fixation, digestion, proximity ligation, and linker sequence enrichment. For detailed protocol, see Supplementary Methods.

In short, the samples were fixed in 30mM HEPES-KOH, 150mM KCl and 1mM DTT by 0.5% formaldehyde and 0.1% glutaraldehyde or 1% formaldehyde. For 96-mer samples, a Kac 12-mer array with independent barcodes was spiked in to estimate the background level at the analysis. The reaction was quenched by glycine, and the samples were dialyzed against the No-salt buffer. The fixed samples were digested by ScaI in the ScaI cut buffer, and 4-16.8 ng of the sample was proximity ligated in 0.125× T4 ligase buffer (50mM Tris-HCl (pH 7.5), 10mM MgCl_2_, 1×mM ATP and 10mM DTT at 1× concentration, New England Biolabs, B0202S) with the presence of 1/250 volume of T4 ligase (Takara Bio, 2011A) at the concentration of 1.05-4.2 pg/µL at 37 °C for 20 min. Proximity ligated samples were deproteinized by adding SDS/PK buffer (10mM Tris-HCl (pH 8.0), 40mM EDTA and 0.5% SDS at final concentration) and Proteinase K (0.5mg/mL at final concentration) and incubating at 65 °C for 2 hours. DNA fragments were concentrated with 2-butanol (Nacalai Tesque Inc., 06102-45), and purified using the FastGene Gel/PCR extraction kit (Nippon Genetics Co., Ltd., FG-91202) with EconoSpin^™^ IIa columns (Ajinomoto Bio-Pharma Services, EP-11201). After excising the linker fragments by NotI-HF (New England Biolabs, R3189S) and 2.5-10U of AvaI, ligated linker fragments were enriched by size selection using Serapure (see Supplementary Methods) and Ampure XP beads.

The sequencing library was prepared from the linker-enriched samples using the NEBNext Ultra II DNA Library Prep Kit for Illumina (New England Biolabs, E7645S) and NEBNext Multiplex Oligos for Illumina (UDI UMI Adaptors DNA Set 1) (New England Biolabs, E7395S) with 13-19cycles of PCR. The PCR cycle was determined for each library using 1/10 of the adaptor-ligated DNA as the template with the KAPA SYBR FAST qPCR Master Mix (Roche, KK4603) by taking the *Ct* that reaches 1/2 of the maximum intensity and subtracting 2 cycles (*Ct* -2 cycles). The entire amount of the remaining adaptor-ligated DNA was used for library amplification. Library pooling was performed ensuring that the linker sequence (observed as a band at 210 bp) was present in equimolar concentration. Sequencing was performed on the NextSeq 2000 sequencer (Illumina) using the P1 flow cell in paired-end mode: 55 cycles for each paired-end read with 20 cycles for index 1 (including 12-base UMI) and 8 cycles for index 2.

### Analysis of AFM data for size measurement

The raw “TOPO” signal recorded for the forward and backward paths was corrected, aligned, and background-subtracted (see Supplementary Methods). The chromatin blobs were found by thresholding the images smoothened by the median filter (3 × 3-pixel footprint) at 0.25 nm, removing objects smaller than 2 pixels and applying the binary closing operation. The histogram for the particle volume (Extended Data Fig. 6b) was smoothed by the Savitzky-Golay filter, and a linear function was extrapolated to the first three peak positions to find the particles with size close to (*n* × 12)-mer with *n* = 1, …, 8. The areas of the particles classified in each *n* were calculated to generate Extended Data Fig. 6c,d.

### Analysis of nanopore sequencing data

To generate the plots in Fig. 1 and Extended Data Figs. 2, the nanopore sequence data was basecalled by the dorado basecaller (version 0.7.2+9ac85c6) [13] with the “sup” model and classified by the barcode using the “demux” command. Sequencing biases were calculated by dividing the number of sequenced base pairs for sequences whose length is in the range [(*n* − 0.5) × *L*, (*n* + 0.5) *L*), where *n* = 1, …, 9 and *L* is the sequence length for a 12-mer array, by the background-subtracted intensity of the electrophoresis intensities in the corresponding bands (Extended Data Fig. 2c).

The linker sequences (barcoded arrays) and the junctions (ligated BglI sites, non-barcoded arrays) were aligned to the reference sequences as described in Supplementary Methods. Using the aligned positions of the linker sequences, we aligned the whole sequences to the entire 96-mer array by offsetting the sequence position by the median of the difference between the expected and sequenced positions (Fig. 1e and Extended Data Fig. 2b). For Fig. 1f and Extended Data Fig. 2h, the histograms were corrected by weighting the reads by the estimated sequence bias (Extended Data Fig. 2c).

For the accessibility assay, the “sup,6mA” model was used to resolve the modified adenine bases. After aligning the sequences to the reference sequences by the “dorado align” command (Supplementary Methods), the frequency of the modified base occurrence was measured using the “pileup” command of modkit (version 0.3.1) [14]. We calculated the accessibility score by subtracting the values for the 0 min samples from those for the 40 min samples and then dividing the values for the chromatin samples by those for the bare DNA samples. The score values smoothed by applying the rolling average of 10 rows were used to plot Fig. 1g and Extended Data Fig. 3.

### Analysis of fluorescence microscopy data

Images were rescaled, preprocessed, and corrected for positional shifts as described in Supplementary Methods. Fluorescent spots were located in the temporally averaged images by using the Laplacian of Gaussian filter (`skimage.feature.blob log` in scikit-image version 0.20.0 [15]) and associated in time (Supplementary Methods). Chromatin molecules that did not fluctuate after compaction or did not elongate with the applied flow were considered non-specifically bound to the coverslip and filtered out by thresholding the Gaussian width (the standard deviation of the fitted Gaussian function) at 50 mM KCl after compaction σ_KCl50 mM_, the ratio of the Gaussian width at 50mM KCl before and after compaction *R*_σ_, and the elongation (distance between the center of the blobs before and after applying the flow) *E*. The blobs with σ_KCl50mM_ > 0.25 µm, 1/1.2 < *R*_σ_ < 1.2, *E* > 0.75 µm were used for further analysis (Extended Data Fig. 5a-c).

28 px 28 × px small movies at the blob positions were exported and further analyzed. First, the images were averaged in time using the sliding window to create images with different exposure times. For each frame, the blob positions were defined as the centroid of the disk-shaped regions around the blobs, with the pixel values weighted by the background-subtracted intensity of the image (Supplementary Methods).

We fitted the median values of the standard deviation of the centroid positions by a theoretical function 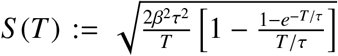, derived as the standard deviation of the positions of a particle following the two-dimensional Ornstein-Uhlenbeck process, 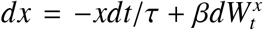 and 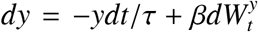, averaged over time *T* corresponding to the exposure time (Fig. 2e and Extended Data Fig. 5e; See Supplementary Methods for derivation).

After fitting, we obtained the standard deviation at the zero exposure limit and the correlation time as 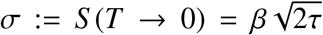 and *τ*, respectively. Considering that we are observing the 2D-projected end positions, we defined the cubed standard deviation of the free end position in Fig. 2f as 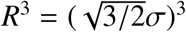.

### Analysis of *in vitro* Hi-C data

The contact maps were generated by locating the ScaI recognition sequence and finding the matching barcode sequences at the expected position (Supplementary Methods). The 12-mer contact maps (Fig. 3c and Extended Data Fig. 9a, extracted from the full contact map) were balanced by the Knight-Ruiz algorithm [16] in the finest resolution resolving all barcode positions. The elements at (*i, i* + *a*) and (*i* + *a, i*) (*a* = 0, 1, 2, 3, *i* = 1, …, *N* − *a, N* is the matrix size) were not used for the balancing, since they are likely to be affected by the ScaI digestion bias. The background level for the 96-mer contact map was calculated by measuring the contact frequency to the spike-in Kac 12-mer array with independent barcodes. The background level was subtracted from the contact map. Stitching was performed for the 96-mer arrays (Fig. 4e-g and Extended Data Fig. 10f,g) by first rescaling the diagonal 12-mer×12-mer submatrices so that they have the same average contact frequency, and then sequentially finding the optimal scale factor for other subcontact maps that minimizes the sum of the squared difference between adjacent regions.

## Supplementary Information

- Supplementary Methods
- Table S1
- References*(1-13)*

## Supplementary Methods

### Plasmid construction for non-barcoded 12-mer nucleosome arrays

To introduce additional restriction enzyme cut sites, the original plasmid containing 12 repeats of the 601 sequence was digested with BbsI-HF (New England Biolabs, R3539), and the plasmid backbone was amplified by polymerase chain reaction (PCR) using the primers containing the sticky end sequences for the ligation reaction. These fragments were then purified by agarose gel electrophoresis and assembled using the NEBuilder HiFi DNA Assembly Master Mix (New England Biolabs, E2621) and transformed into NEB 5-alpha Competent *E. coli* (New England Biolabs, C2987).

### Plasmid construction for barcoded 12-mer nucleosome arrays

First, a plasmid with a single 601 sequence and spacer sequence (plasmid 1) was generated by PCR amplification of the plasmid backbone of a non-barcoded plasmid, digesting the product by AflIII (New England Biolabs, R0541S) and BanI (New England Biolabs, R0118S), which was ligated to a single 601 fragment digested by AflIII and BanI. The plasmid was then amplified using the 5-alpha cells and purified using the NucleoSpin^™^ Plasmid Transfection-grade kit (Macherey-Nagel, 740490), digested with AvaI (New England Biolabs, R0152S) and purified by agarose gel electrophoresis.

Fragments containing a 601 sequence and a barcoded linker sequence were generated by annealing two oligonucleotides (120 bp each, 20 bp annealing region) with a temperature gradient from 95 to 4 °C at 1 °C/min in buffer containing 10mM Tris-HCl (pH 7.5) and 50mM NaCl or 10mM Bis-Tris-Propane (pH 7.4) and 100mM NaCl, and then filling in the single stranded parts by 0.1U Klenow fragments (Takara Bio, 2140A) at the concentration 0.1 µg/µL with 0.1mM dNTP mix (New England Biolabs, N0447S) in 1X Klenow fragment buffer (10mM Tris-HCl (pH 7.5), 7mM MgCl_2_, 0.1mM DTT). The sample was incubated first at 10 °C for 4 hours and then at 4 °C overnight, and then precipitated by ethanol with an overnight incubation at − 20 °C. 40 µg of the filled oligonucleotide was cut with 200U AvaI in 200 µL and purified by electrophoresis in 3% agarose gel.

The digested fragment was then ligated into the AvaI-digested plasmid 1 treated with Quick CIP, transformed into the 5-alpha cells, and the cells were grown on an LB plate. The colonies were then collected and midi-cultured, and the plasmid was purified using the NucleoBond Xtra Midi kit (Macherey-Nagel, U0410B) (plasmid 2).

Fragments containing a single 601 sequence and a barcoded linker sequence with the BsmBI recognition sequence cutting the AvaI site were similarly prepared and ligated into the BbsI-digested plasmid 2 treated with Quick CIP (BbsI-HF: New England Biolabs, R3539L). The plasmid was transformed into the 5-alpha cells, which were grown on an LB plate. Colonies were collected, and the plasmids were purified using the NucleoSpin Plasmid Transfection-grade kit (plasmid 3)

Fragments containing a 601 sequence and a barcoded linker sequence were purified from the plasmid 2 by digestion with AvaI and electrophoresis in 3% agarose gel. 180 ng of the fragment was then ligated to the plasmid 3 (digested by BbsI and treated with Quick CIP)at a 10:1 ratio with 2000U T4 ligase (New England Biolabs, M0202M), 16 °C for overnight in 13 µL. The ligated product was transformed into the 5-alpha cells, which were grown on an LB plate. The colonies were then collected and midi-cultured, and the plasmid was purified using the NucleoBond Xtra Midi/Maxi kit (plasmid 4). The plasmid 4 was then transformed into the 5-alpha cells, which were grown on an LB plate. The colonies were isolated and checked by direct PCR, which revealed that the plasmid typically contains 4–9 repeats of the 601 sequence along with the spacer sequences.

Colonies hosting plasmids containing 5-mer to 7-mer repeats without errors were identified by direct colony PCR and Sanger sequencing. Plasmids were isolated from those colonies using the NucleoSpin Plasmid Transfection-grade kit, digested by BbsI, and treated with Quick CIP. Nine independent plasmids were then constructed by ligating the digested plasmids to the 5-mer to 7-mer products isolated from plasmid 4 using BbsI and BsmBI digestion (BsmBI-v2: New England Biolabs, R0739S) followed by agarose gel electrophoresis. The BglI sites for the sticky ends and the adjacent barcode sequences were modified.

### Purification of octamer sink DNA fragments

2-300 bp octamer sink DNA fragments were generated by taking the supernatant of the PEG precipitation to concentrate the EcoRV-digested backbone fragments, precipitating the DNA with ethanol, and digesting the remaining 12×601 array with StyI-HF (New England Biolabs, R3500S) and precipitating by phenolchloroform-isoamyl alcohol and ethanol if it remained in the product. These octamer sink fragments were used in the reconstitution step to prevent non-specific binding of histone octamers to linker regions [1, 2].

### End labeling of A and H fragments

For labeling, 28 µL of 1mM modified oligonucleotide (5’-Ph-TAGTCTGC(T*)CAGATATCGTCG-Biotin-3’, where T* represents the amine-linked dT with the C6 linker, purchased from Merck) dissolved in 0.091 M sodium tetraborate-HCl buffer (NaB buffer, made from Sodium Tetraborate Decahydrate, Nacalai Tesque, Inc., 31223-85) was mixed with 5.6 µL of 14mM Atto647N-NHS ester (ATTO-TEC GmbH, AD647N-31) in DMSO and incubated overnight at room temperature. The labeled fragment was ethanol precipitated with ×1/3.5 volume of 3M NaOAc and ×9/3.5 volume of ethanol, with 10 min incubation at 20 °C. Another modified oligonucleotide (5’-CACAGCTG(T*)CTGAG-3’) was also labeled with Atto565-NHS ester (ATTO-TEC GmbH, AD 565-31). The dye labeling efficiency measured by the absorbance was 60% and 69%, respectively.

These labeled oligonucleotides were then annealed with the complementary strands extended by 3 bp for the sticky ends that ligate with the A and H arrays, respectively (Fig. 1b). Annealing was performed at a concentration of 100 µM for each fragment, in the presence of 10mM Bis-Tris-Propane (pH 7.4) (Nacalai Tesque, Inc., 08892-32), 1mM MgCl_2_ and 100mM NaCl, with cooling at a rate of 1 °C/min.

1.2 mg of the BglI-digested and PEG-purified A and H fragments were then mixed with a five-fold molar excess of the annealed products and ligated with 2100U T4 ligase (Takara Bio, 2011A) in a final reaction volume of 750 µL at 16 °C for overnight. The ligated product was then PEG-purified as described in the previous section.

### Reconstitution of 12-mer nucleosome arrays

The template DNA hosting 12 repeats of the 601 sequence was mixed with the reconstituted histone octamer and short octamer sink DNA fragments, where the salt concentration was adjusted to 2M KCl. The molar ratio of the octamer and buffer DNA fragment to the 601 sequences was chosen to be 1.2 and 5/12 for the non-barcoded arrays and 1.1 and 1/3 for the barcoded arrays.

The mixture was transferred to a microdialysis tube, Xpress Micro Dialyzer MD100 MWCO 6-8 kDa (Scienova GmbH, SCI-40076), and dialyzed in a gradient from RB-High (10mM Tris-HCl (pH 7.5) (Nacalai Tesque, Inc., 35434-21), 1mM EDTA (Nacalai Tesque, Inc., 06894-85), 2M KCl (Nacalai Tesque, Inc., 28514-75), and 1mM DTT (Nacalai Tesque, Inc., 14128-62)) to RB-Low (10mM Tris-HCl (pH 7.5), 1mM EDTA, 250mM KCl) by adding 1.6 L of RB-Low to 0.4 L of RB-High at a rate of 1.6mL/min while maintaining the total volume. The dialysis tubes were then transferred to 400mL of RB-Low for another 4 hours of dialysis. The mixture was transferred to 1.5mL microcentrifuge tubes and centrifuged at 10, 000 × *g* for 10 min. The supernatant was again transferred to microdialysis tubes and dialyzed against 400mL of No-salt buffer (10mM Tris-HCl (pH 7.5) and 1mM EDTA) for more than 4 hours. The mixture was then transferred to 1.5mL microcentrifuge tubes and centrifuged at 10, 000×*g* for 10 min, and the supernatant was transferred to fresh lo-bind tubes (BMBio ST, BM4015).

After the reconstitution, the arrays were purified by 10% - 40% sucrose gradient ultracentrifugation. The centrifuged solution was fractionated into 800 µL aliquots by gentle aspiration from the top of the tube, and each fraction was checked by the agarose gel electrophoresis (0.7% agarose in 0.2X TBE). One or two peak fractions were then concentrated using 100 kDa MWCO Amicon Ultra centrifugal filter (Merck Millipore, UFC5100) at 14, 000 × *g*, and then washed twice by adding 450 µL of No-salt buffer and centrifuging at 14, 000 × *g* for 5 − 15 min.

Reconstituted chromatin was routinely checked by the band shift assay in agarose gel electrophoresis (0.7% agarose in 0.2X TBE). Also, the nucleosome reconstitution efficiency was validated by digesting ≈ 100 ng of the sample with 2.5 − 10U of ScaI (Takara Bio, 1084A) in 10 µL of the ScaI cut buffer (10mM Tris-HCl (pH 7.5), 0.5mM MgCl_2_ and 50mM NaCl) at 22 °C for 14 hours and confirming the band shift with electrophoresis in native polyacrylamide gel electrophoresis (PAGE) (5% in 1X TBE, 120V for 30 min) (Extended Data Fig. 1).

The reconstituted 12-mer arrays were diluted to 50-100 ng/µL and supplemented with glycerol at a final concentration of 5%, aliquoted to 2.5 µg per tube, snap frozen with liquid nitrogen, and stored at −80 °C.

### Details of AFM observation

#### Sample fixation

For the nucleosome counting (Fig. 1 and Extended Data Figs. 1, 4, except Fig. 1i and Extended Data Figs. 4a,e-m), samples were diluted in the No-salt buffer to 0.25-1 ng/µL and fixed with 0.1% glutaraldehyde (Nacalai Tesque Inc., 17003-92) on ice for 10 min (12-mers) or 30 min (96-mers). For Fig. 1i and Extended Data Figs. 4a,e-m, the sample was diluted in H30D buffer (30mM HEPES-KOH (pH 7.5) and 1mM DTT) to 3 ng/µL and fixed with 0.1% formaldehyde (FUJIFILM Wako Pure Chemical Corp., 064-00406) and 0.05% glutaraldehyde on ice for 15 min.

#### Mounting sample on mica plate for AFM observation

A mica plate (Alliance Biosystems, 01872-MB) was peeled by a piece of adhesive tape. 0.01% poly-L-ornithine solution (Sigma-Aldrich, P3655) was spotted onto the freshly peeled mica plate and then incubated for 1.5 min at room temperature. The plate was then washed three times with 1mL of ultrapure water (Milli-Q, Merck Millipore) and dried completely with a stream of nitrogen gas.

For the 12-mer samples and 96-mer samples except Fig. 1h,i and Extended Data Figs. 4a,e-m, sample solutions were spotted onto the coated mica plate and incubated for 5 min at room temperature. The sample was gently washed out with 1mL of ultrapure water and dried with a stream of nitrogen gas. For Fig. 1h, a 1.5mm × 3mm chamber (1mm thickness) made of PDMS sheet and a glass slide was placed on the coated mica plate and 50 µL of the sample was gently injected in 10 sec. We then removed the chamber, washed the plates with 1mL of ultrapure water twice, and dried them with a stream of nitrogen gas. For Fig. 1i and Extended Data Figs. 4a,e-m, 10 µL of the sample was loaded onto the mica plate spinning at 850-1000 rpm for 20 sec. After the spin-coating, the sample was gently washed out with 1mL of ultrapure water and dried with a stream of nitrogen gas.

Samples were observed with the Innova atomic force microscope (Bruker) using the SNL10-C probe in the tapping mode. The set point was typically at 1.5-2V with the drive amplitude at 5 V.

### Details of single-molecule fluorescence microscopy

#### Chamber construction for single-molecule observation

For the chamber construction for single-molecule imaging, we followed a protocol in Ref. [2] with modifications. Inlet and outlet holes were made on glass slides (Matsunami Glass, S2441) with a micro-grinder. Coverslips (Matsunami Glass, No.1SHT) and glass slides were placed on a Teflon rack in a glass beaker. They were washed by sonication in 20× diluted Ultrasonic Cleaner Alkaline Cleaning Agent (SDNU-A4, AS ONE Corporation) for 60 min by ASU-2 (AS ONE Corporation) and kept overnight. The glasses were rinsed five times with ultrapure water and sonicated for 20 min in ultrapure water. They were then rinsed with acetone, sonicated for 20 min in acetone, rinsed with isopropanol, sonicated for 20 min in isopropanol, rinsed with ultrapure water five times, sonicated for 20 min in ultrapure water, and rinsed with ultrapure water five times. The coverslips were Piranha-cleaned with 25% v/v Hydrogen Peroxide (30%) (Nacalai Tesque, Inc., 18411-25) and 75% v/v Sulfuric Acid (Nacalai Tesque, Inc., 32519-95) at 80 °C for 60 min and rinsed four times with ultrapure water. Then, they were mildly etched with 0.5 M KOH with sonication for 60 min to increase the density of the silanol group. The sonication chamber (14 cm×15 cm×10 cm) was first filled with ice to avoid heating [3]. The glasses were rinsed twice with ultrapure water and then twice with acetone. The surface was amino-activated by soaking in either 100mL of 3% 3-aminopropyltriethoxysilane (Nacalai Tesque, Inc., 02309-04) in the mixture of 95mL of ethanol, 5mL of ultrapure water, and 2mL of acetic acid (Replicate 1) or 80mL of 3% 3-aminopropyltriethoxysilane in acetone (Replicate 2) for 60 min at room temperature. The glasses were then rinsed twice with acetone, then twice with ultrapure water, and dried with a stream of nitrogen gas.

40 mg of PEG-SVA (MPEG-SVA, Laysan Bio, Inc.) and 1 mg of biotin-PEG-SVA (BIO-PEG-SVA, Laysan Bio, Inc.) were dissolved into 120 µL of NaB buffer with 0.1 M K_2_SO_4_. The solution was vortexed well and centrifuged at 10, 000 × *g* for 15 sec. Immediately, 40 µL of the solution was dropped onto the amino-modified coverslip, and another coverslip was placed to sandwich the solution. The coverslips were kept overnight at room temperature in a dark, humid chamber and then quickly rinsed with ultrapure water and dried with a stream of nitrogen gas. They were further passivated with short PEG-NHS ester (2,5-dioxopyrrolidin-1-yl 2,5,8,11-tetraoxatetradecan-14-oate, BLD, BD306356). 12 µL of short PEG was mixed with 240 µL of NaB buffer containing 0.5M KCl, and the coverslips were reacted with the solution in the same way for 2 hours at room temperature. After the reaction, they were quickly rinsed with ultrapure water and dried with a stream of nitrogen gas. For replicate 1, the surface was further passivated by incubation with 1 mg of *N*-Succinimidyl Acetate (Tokyo Chemical Industry, S0878) dissolved into 240 µL of NaB buffer containing 0.5 M KCl, for 2 hours at room temperature. After the reaction, they were quickly rinsed with ultrapure water and dried with a stream of nitrogen gas.

Pieces of double-coated Kapton^®^ film tape (silicone-based adhesive, Teraoka 760H), ≈ 45 µm thick, were cut to make five 12 mm × 2 mm channels. SYLGARD^™^ 184 silicone elastomer kit (DOW) was used to fabricate a polydimethylsiloxane (PDMS) block. 20 g of base and 2 g of hardener were mixed well, degassed in a vacuum chamber, poured into a 10 cm diameter petri dish, and cured overnight at 80 °C. PDMS blocks were cut out and holes punched to match the holes in the glass slide. The glass slide and PDMS mold were then plasma etched (“Hi” configuration in PDC-32G, Harrick Plasma, corresponding to 18 W applied power to the RF coil) for 1 min and immediately pressed together for adhesion.

The chamber was constructed by sandwiching the adhesive tape between the glass slide and passivated coverslip (Fig. 2b). The channel height was measured with a confocal microscope and found to be 57 ± 6 µm. The chambers were kept in a vacuum at − 20 °C.

At observation, the chambers were taken out of the freezer, and liquid reservoirs (1mL pipette tips) were inserted into the holes of the PDMS mold and glued with epoxy. We waited for more than 15 min to avoid contaminating the observation chambers with the glue. The coverslips were further passivated again by injecting 20 µL of the mixture of short PEG-NHS ester (5 µL) and NaB buffer (100 µL) (replicate 1) or 0.1M NaB buffer (pH 8.5) containing 0.5M KCl (replicate 2) and incubating for more than 15 min.

#### Imaging buffer preparation

37.5 mg of (±)-6-Hydroxy-2,5,7,8-tetramethylchromane-2-carboxylic acid (Trolox, Sigma-Aldrich, 238813) was mixed with 37.5mL of ultrapure water supplemented with 50 µL of 2.8M NaOH. The solution was rotated in the dark until the precipitate dissolved, and the concentration of Trolox was measured by the absorbance at 290 nm (2350M^−1^cm^−1^). The Trolox solution was then supplemented with HEPES-KOH (pH 7.5, Dojindo 348-01372), D(+)-glucose (FUJIFILM Wako Pure Chemical Corp., 045-31162), and KCl to make the IBs containing 50mM HEPES-KOH (pH 7.5), 2mM Trolox, 3.2% w/v glucose and 50mM, 100mM or 150mM of KCl. Glucose oxidase + bovin liver catalase solution (GODCAT) was prepared by first dissolving 1 mg of catalase (C40-100MG, Sigma Aldrich) in 100 µL of 50mM PBS (pH 7), and mixing the 20 µL of this solution with 30 µL of T50 (10mM Tris-HCl (pH 7.5) and 50mM NaCl) and 5mg of glucose oxidase (Sigma Aldrich, G2133-10KU). The IBs were supplemented with 1/200 volume of GODCAT just before observation.

#### Microscope setup

The sample was observed with a fluorescence microscope built with IX83 (Olympus) equipped with the objective lens UPlanApo 60x / 1.50 Oil HR TIRF (Olympus) and with the filters and dichroic mirrors of ZET405/488/561/640xv2, ZT405/488/561/640pcv2 and ZET405/488/561/640mv2 (Chroma Technology). The laser light from OBIS 561nm LS (Coherent) and OBIS 637nm LX (Coherent) were combined by a dichroic mirror (ZT561rdc-UF3, Chroma Technology) and introduced from the rear port of the microscope and focused at the rear focal plane of the objective.

The Multisplit V2 (Teledyne Photonics) split the fluorescence light into regions. Fluorescent light that

- Passed through T565lpxr and T635lpxr, and filtered by ET655lp (Chroma Technology)
- Passed through T565lpxr and reflected by T635lpxr, and filtered by ET600/50m (Chroma Technology)

were observed as the signals for the tethered end (Atto647N) and free end (Atto565), respectively (Fig. 2a,c). The angle of the laser has been adjusted for observation with the Highly Inclined and Laminated Optical Sheet (HILO) microscopy [4] so that the signal from the Atto565 is maximized. The laser intensity was 10-20mW (561 nm) and 10mW (640 nm), respectively.

### Detailed protocol for *in vitro* Hi-C experiment

#### Fixation

Samples were dialyzed against 12mL of H30D buffer (30mM HEPES-KOH (pH 7.5) and 1mM DTT) for 1 hour at 4 °C in the microdialysis tubes before fixation. They were fixed at the final concentration of 4 ng/µL (12-mer samples), 2.37 ng/µL (96-mer samples, replicates 1 and 2) or 2 ng/µL (96-mer samples, replicates 3 and 4) with 0.5% formaldehyde and 0.1% glutaraldehyde in 30mM HEPES-KOH, 150mM KCl and 1mM DTT at 30 °C for 5 min. The reaction was quenched by adding×1/11 volume of 12×quenching buffer (2.12 M glycine (Nacalai Tesque Inc., 17141-95) and 120mM Tris-HCl (pH7.5, Nacalai Tesque Inc., 35434-21 and 37313-25)). For the 12-mer experiment, the fixation and quenching were performed with the combination of 1% formaldehyde and ×1/6 volume of the quenching buffer 2 (2.33 M Glycine and 70mM Tris-HCl (pH 7.5)) and 0.5% formaldehyde + 0.1% glutaraldehyde and ×1/12.21 volume of the quenching buffer 2. Samples were dialyzed against 12mL of No-salt buffer for 1 hour at 4 °C in the microdialysis tubes.

The sample after fixation was checked by the agarose gel electrophoresis and found to show a similar band pattern to that for the samples without fixation (Extended Data Fig. 8a,b). The sample concentration was measured using the Qubit^™^ dsDNA Quantification Assay Kit HS (Invitrogen, Q32854), and the measured value was used to dilute the sample in the following steps. The measured value was ≈ 2.1 times lower than the actual DNA concentration due to the lower dye binding efficiency. In the following steps, the concentration was calculated by multiplying the measured value by 2.1.

#### ScaI digestion and wash

The samples were digested at the center of the linker with 0.5 U/µL ScaI (Takara Bio, 1084A) in the ScaI cut buffer (10mM Tris-HCl (pH 7.5), 0.5mM MgCl_2_, 50mM NaCl, and 1mM DTT supplemented with 0.1 mg/mL of recombinant albumin (New England Biolabs, B9200S)) at 22 °C for 14 hours at a final concentration of 5. 6-6.1 ng/µL (12-mer samples), 3.6 ng/µL (96-mer samples, replicates 1 and 2), or 3.0 ng/µL (96-mer samples, replicates 3 and 4). Unfixed samples were also digested at a similar concentration.

The sample was mixed with ×1/9 volume of 10× Mg^2+^ quench buffer (500mM Tris-HCl (pH 7.5) and 190mM EDTA) and incubated at 22 °C for 20 min. 5.25-12.1 ng of the sample was taken and digested by SDS/PK (0.5 mg/mL Proteinase K 10mM Tris-HCl (pH 8.0), 40mM EDTA and 0.5% SDS) at 65 °C for 2 h and checked by PAGE and Agilent 2100 Bioanalyzer High Sensitivity DNA Kit (Agilent, 5067-4626) (Extended Data Fig. 8c-g). The results confirm that the DNA was digested into mononucleosomes without systematic bias in the digestion efficiency dependent on the histone modification (Extended Data Fig. 8f,g).

The samples were washed using 100 kDa MWCO Amicon Ultra centrifugal filters using the following protocol:

1. Load 500 µL of the Chromatin Wash Buffer (20mM HEPES-NaOH (pH7.5), 50mM NaCl, 0.5mM EDTA and 0.05% IGEPAL CA-630 (MP Biomedicals, Inc., 198596)) and centrifuge at 6000 × *g* for 3min at 4 °C.
2. Load the samples and 400 µL of the Chromatin Wash Buffer. Centrifuge at 6000 × *g* for 3min at 4 °C.
3. Add 450 µL of the Chromatin Wash Buffer, and centrifuge at 6000 × *g* for 3min at 4 °C.
4. Add 450 µL of the Chromatin Wash Buffer, and centrifuge at 6000 × *g* for 5min at 4 °C.
5. Add 450 µL of the No-salt buffer and centrifuge at 6000 × *g* for 5min at 4 °C.

The washed samples were quantified using the Qubit HS kit. 1.1-10.5 ng of the samples were used for quantification by PAGE (Extended Data Fig. 8c,d). The results indicate the appearance of multi-nucleosome bands, which were not observed in the unfixed samples (Extended Data Fig. 8c,d).

#### Proximity ligation

Proximity ligation of 4-16.8 ng of the sample was performed in 0.125× T4 ligase buffer (50mM Tris-HCl (pH 7.5), 10mM MgCl_2_, 1mM ATP and 10mM DTT at 1× concentration, New England Biolabs, B0202S) with the presence of 1/250 volume of T4 ligase (Takara Bio, 2011A) at the concentration of 4.2 pg/µL (12-mer samples and 96-mer sample replicate 1) and 1.05 pg/µL (other samples) at 37 °C for 20 min. To avoid aggregation, we first diluted the samples in 10mM Tris-HCl (pH 7.5), and then ligase buffer and T4 ligase were added and mixed by inversion of the tubes (PROTEOSAVE SS 15mL Conicaltube, FUJIFILM Wako Pure Chemical Corp., 634-28101).

After the proximity ligation, the samples were deproteinized by adding SDS/PK buffer (10mM Tris-HCl (pH 8.0), 40mM EDTA and 0.5% SDS at final concentration) and Proteinase K (0.5mg/mL at final concentration) in ×1.4 volumes of the proximity ligation and incubating at 65 °C for 2 hours. The samples were concentrated with 2-butanol (Nacalai Tesque Inc., 06102-45) to 3-600 µL, and purified using the FastGene Gel/PCR extraction kit (Nippon Genetics Co., Ltd., FG-91202) with EconoSpin^™^ IIa columns (Ajinomoto Bio-Pharma Services, EP-11201). ×5 volume of the GP1 solution was used in the first step. The final product was eluted to 40 µL of GP3, which was pre-warmed to 60 °C.

The concentrated samples were quantified using the Qubit HS kit. 0.5 ng of the samples were used for quantification by Agilent 2100 Bioanalyzer High Sensitivity DNA Kit (Extended Data Fig. 8e). The result indicates the appearance of the ligation product compared to the input samples.

#### Linker digestion, enrichment, and library preparation

The linker sequences were excised from the ligated product by digestion using 32 µL (20 µL for the 12-mer samples) of sample, 5-20U of NotI-HF (New England Biolabs, R3189S) and 2.5-10U of AvaI in 1×rCutSmart buffer (New England Biolabs), 37 °C (lid 75 °C) for 2 hours in 40 µL (25 µL for the 12-mer samples). The digested product was quantified by Agilent 2100 Bioanalyzer High Sensitivity DNA Kit (Extended Data Fig. 8e).

Magnetic bead solution for purification [5] (Serapure) with 2.5M NaCl was prepared according to the following protocol:

1. Take 1mL of Sera-Mag SpeedBead Carboxylate-Modified Magnetic Particles (Cytiva, 65152105050250) and wash them with 1mL of 10mM Tris-HCl (pH 8.0) (Nacalai Tesque Inc., 35435-11) by pipetting three times. Place the solution on a magnetic rack and discard the supernatant.
2. Add 1mL of 100mM Tris-HCl (pH 8.0) to the washed beads and mix well.
3. Add 25mL of 5M NaCl (Nacalai Tesque Inc., 31320-05) and the bead solution into a 50mL conical tube.
4. Add 20mL of 50% (w/v) PEG 8000 (MP Biomedicals, Inc., 195445) in ultrapure water to the tube.
5. Add 4mL of 100mM Tris-HCl (pH 8.0) and adjust the final volume to 50mL by adding ultrapure water.

The sample was purified, and short linker fragments were enriched using Serapure, Ampure XP beads, and precipitation solution (PS, 2.5M NaCl, 20% PEG 8000, 10mM Tris-HCl (pH 8.0)) according to the following protocol:

1. Pre-enrichment wash
  (a) Add 120 µL of Serapure:PS = 1.8:1.2mix to 40 µL of sample, pipette 15 times.
  (b) Incubate for 5 min at room temperature.
  (c) Place the sample on a magnetic rack and wait for 2 min.
  (d) Discard the supernatant.
  (e) Add 100 µL of 70% ethanol and discard.
  (f) Repeat the previous step.
  (g) Remove the sample from the magnetic rack, centrifuge, and place it on the magnetic rack. Remove the ethanol and dry the beads briefly, but do not wait longer than 1 min.
  (h) Add 10 µL of TE buffer and pipette well. Wait for 2 min.
  (i) Place the sample on a magnetic rack and take the supernatant to another PCR tube.
2. Linker enrichment
  (a) Add 10 µL of TE buffer to the sample. Mix the sample, Ampure XP Beads, and PS in a ratio of 1:1.2:0.3 (20 µL, 24 µL and 6 µL) by pipetting 15 times.
  (b) Place the samples on the ThermoMixier C (Eppendorf) and keep 25 °C 600 rpm and incubate for 30 min.
  (c) Immediately place the samples on a magnetic rack and incubate for 2 min.
  (d) Transfer the supernatant to another PCR tube.
  (e) Add 150 µL of Serapure and PS solution (ratio 2:1) to the supernatant. Mix by pipetting 15 times.
  (f) Incubate the samples at room temperature for 5 min.
  (g) Place the samples on a magnetic rack and wait for 2 min.
  (h) Discard the supernatant.
  (i) Add 100 µL of 70% ethanol to the beads and discard. Repeat this step twice.
  (j) Remove the sample from the magnetic rack, spin it quickly, and place it on the magnetic rack to remove all ethanol. Dry the beads briefly, but do not wait longer than 1 min.
  (k) Add 5uL TE and pipette well. Wait 2 min at room temperature.
  (l) Place the samples on a magnetic rack and transfer the supernatant to another PCR tube.

The samples were analyzed using the Agilent 2100 Bioanalyzer High Sensitivity DNA kit (Extended Data Fig. 8e). The result indicates the enrichment of the linker fragments compared to the input samples.

The sequencing library was prepared from the linker-enriched samples and sequenced as described in the main text.

#### Preprocessing for AFM images

The forward and backward “TOPO” signals from the AFM were opened with Gwyddion [6] (version 2.62) and processed with the functions “Correct scars”, “Align rows” (in the “Matching” mode), “Level image” and then saved as HDF5 files. Row-wise backgrounds for these signals were estimated by taking the column-wise median values of the pixels, excluding the pixels that deviated more than 30 nm from the median height for the row. The signals were rescaled by subtracting this row-wise background.

The column offset to align the forward and backward signals was found by the peak of the cross-correlation of the two signals, and these signals were merged taking the minimum value of the aligned images similarly to [7]. After that, the two-dimensional background signal was estimated as follows. First, the median filter with a footprint of 5 × 5 was applied, followed by applying a Gaussian filter with a standard deviation of 2. The background was then estimated using a rolling ball algorithm (`skimage.restoration.rolling ball`) with a radius of 10 after scaling the image by a factor of 2 ×10^11^, followed by an additional Gaussian filter with a standard deviation of 5. The result was then rescaled to the original range by dividing by the scaling factor. We performed another round of background subtraction by estimating the background height using the `Baseline2D.pspline arpls` function from the pybaselines package [8] (version 1.1.0). The mask image for the pixels used for fitting was generated by thresholding the image with 0.25 nm and eroding with a 4-pixel radius disk.

### Details of nanopore sequencing data analysis

#### Alignment of linker sequences

For the barcoded arrays, the linker sequences were aligned to the reads using `Bio.Align. PairwiseAligner` of Biopython (version 1.83) [9] with the settings

- `open gap score = -2`
- `extend gap score = -4`
- `query left open gap score = -10`
- `query left extend gap score = -10`
- `query right open gap score = -10`
- `query right extend gap score = -10`
- `mode = ‘local’`

The alignment results with an alignment score greater than 42 and mapped to a region containing 5-55 bp of the query were analyzed downstream. Since the length of the aligned sequences was different for the 5’-end and 3’-end of the 96-mer, the threshold score was calculated as 0.8 × *l*, where *l* is the length of the aligned sequence. F or the non-barcoded arrays, the junctions (ligated BglI sites) were searched using the `Bio.SeqUtils.nt search` function.

#### Alignment for accessibility assay

First, the linker sequences were aligned as described. Sequences were categorized by the containing 12-mers and exported into BAM files, retaining the information from the base-called BAM files. The sequences in the BAM files were then aligned to the corresponding reference sequence by the “dorado align” command.

### Details of fluorescence microscopy data analysis

#### Preprocessing and molecule finding

Image data were rescaled by subtracting the background signal (measured from the mean signals without incident light) and dividing by the mean pixel gain (measured by fitting a linear function to the mean and standard deviation values at low illumination). Images for each condition (100 frames each) were averaged for blob detection. Blobs were detected using the Laplacian of Gaussian filter (`skimage.feature.blob log` in scikit-image version 0.20.0 [10]) with the parameters `min sigma=2, max sigma=2, num sigma=2, threshold=0.75, overlap=0.25, exclude border=25`. For each blob, two-dimensional Gaussian functions were fitted to the data with the objective of the least square difference by successively applying the Nelder-Mead method and the trust-constr method using the symfit package [11] (Version 0.5.6). Blobs with a Pearson correlation of less than 0.5 from the fitted function values, less than 50 photons for a single frame, and whose center deviates from that detected by the Laplacian of Gaussian filter were excluded from the analysis.

The positional shift between the two channels was corrected using images of multicolor fluorescent beads (TetraSpeck Microspheres, 0.2 µm, Thermofisher, T7280). The movement of the stage between conditions was corrected using the positions of the far-red blobs (the tethered ends). Specifically, the translation and rotation that minimizes the cost function 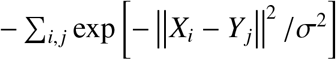, where *X*_*i*_ and *Y*_*j*_ are the blob positions before and after the transformation, were found by the brute-force method. Then, the blobs without a matching blob within 2 pixels were discarded, and the final translation and rotation values were estimated by minimizing the mean squared Euclidean distance to the nearest blob. We tracked the blob positions for the free ends in the aligned coordinate to associate them temporally. Since the blob typically shifts with the flow, the images for the flow-applied conditions were shifted to compensate for the effect of the flow (for this association process only). We used LapTrack [12] with `track dist metric=“sqeuclidean” and track cost cutoff=225`. The blobs not tracked for more than 50 frames were removed from the analysis. Free ends within 3 pixels of the nearest tethered end were classified as colocalized.

#### Blob position analysis

For each image, the images were thresholded at the 90% percentile of intensity, and connected regions with less than two pixels and those touching the boundaries were removed. Movies with more than two connected regions after dilation with the kernel of a 3-pixel radius disk and those with more than five pixels in the thresholded regions touching the boundaries were removed from the analysis. The “centroid masks” were created as disk-shaped regions with a radius of 5 pixels whose center was at the centroid of the original segmented region. The blob position is defined as the centroid of the centroid mask, with the pixel values weighted by the background-subtracted intensity of the image.

#### Derivation of the fitting function for different exposure times

We obtained the formula 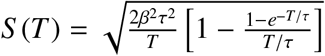 by calculating the variance of the time-averaged position 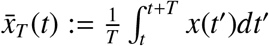 as

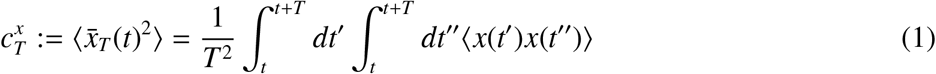

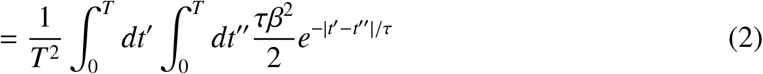

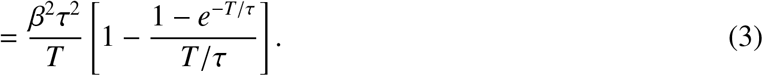

The standard deviation of the free end position in 2D (Fig. 2e and Extended Data Fig. 5e) is therefore given by 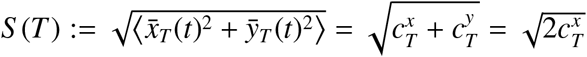.

#### Zimm model with tethered end

Consider a polymer chain consisting of *N* beads connected by harmonic springs. The time evolution of the position of the *i*-th bead, 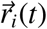, is governed by the Langevin equation

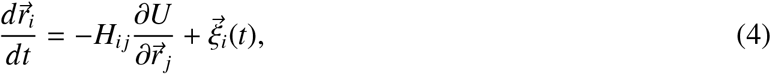

where *H* is the hydrodynamic interaction tensor, *U* is the potential energy of the system, and 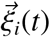 is the random force acting on the *i*-th bead (*i* = 0, 1, …, *N* − 1). The random force satisfies the fluctuation-dissipation theorem

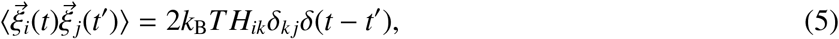

where *k*_B_ is the Boltzmann constant, *T* is the temperature, and⟨·⟩ denotes the ensemble average. Let us consider the Rouse model, where the potential energy is given by

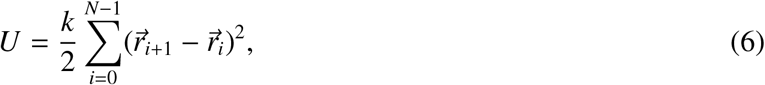

where *k* is the spring constant that satisfies

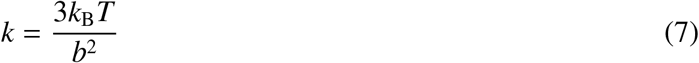

with *b* being the Kuhn length. Under this setup, the average end-to-end distance of the polymer is 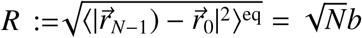, where ⟨·⟩_eq_ denotes the equilibrium ensemble average.

In the Zimm model, the hydrodynamic interaction tensor is substituted by its equilibrium value:

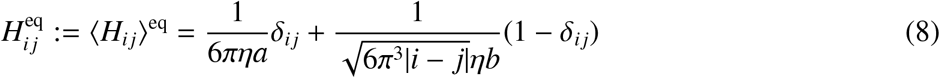

where *η* is the viscosity of the solvent, and *a* is the radius of the bead. The Rouse model is obtained by neglecting the off-diagonal terms in the hydrodynamic interaction tensor, which is justified if *a* ≪ *b*.

The Langevin equation is now given by

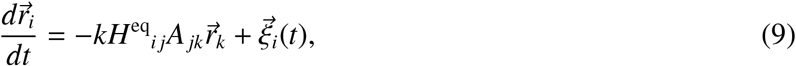

Where

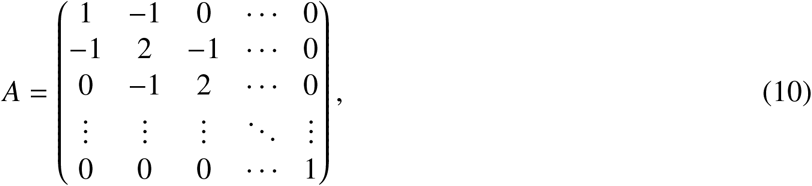

for the case of free ends, and

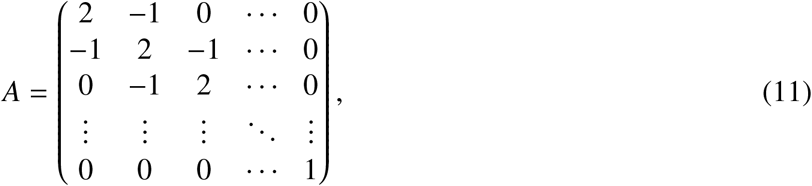

for the case of one end tethered to a fixed point. We note that the effect of the impenetrable interface at the tethered end (e.g., passivated glass) is negligible since the experiment measures fluctuations parallel to it. In this case, the relaxation time scale is identical to the chain tethered at one end but otherwise free [13].

By diagonalizing the matrix *A* as *A* = *O*Λ*O*^*T*^ using an orthogonal matrix *O* and a diagonal matrix Λ, the equation of motion can be written as

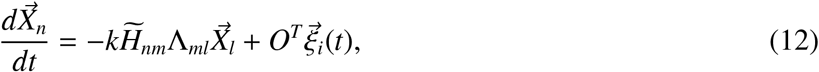

with 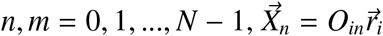 and 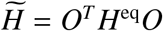.

For the free end case, the eigenvalues of *A* are (for *n* = 0, 1, …, *N* − 1)

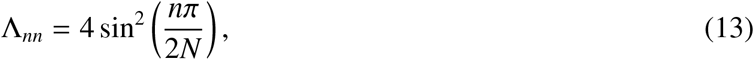

and the eigenvectors are given by (for *i* = 0, 1, …, *N* − 1 and *n* = 0, 1, …, *N* − 1)

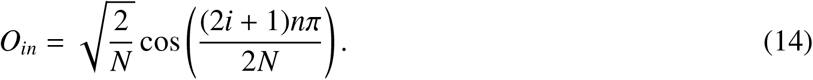

We therefore have (for *n, m* = 0, 1, …, *N* − 1)

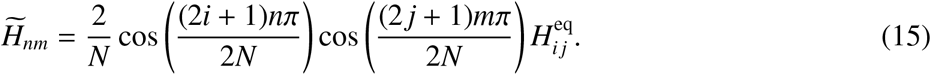

For the tethered end case, the eigenvalues of *A* are

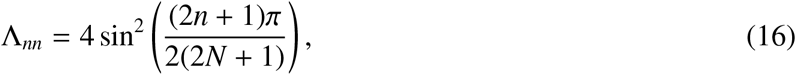

and the eigenvectors are given by

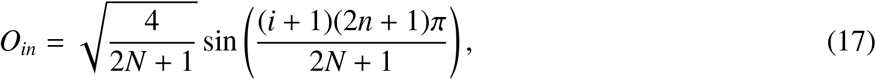

We therefore have

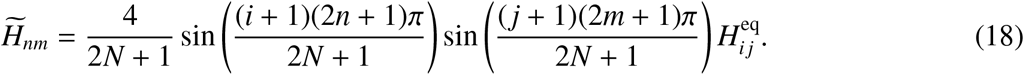

The slowest time scale of relaxation is given by the inverse of the smallest non-zero eigenvalue of 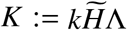. However, this is, in general, an ill-defined problem, since the eigenvalues of *K* can be negative, when the off-diagonal part of the hydrodynamic interaction tensor becomes dominant. This unphysical situation arises due to the approximation Eq. (8). We nevertheless expect that this approximation is valid for the spatially extended modes (i.e., small *n*), and take the eigenvalue corresponding to the eigenvector that has the largest component of the *n* = 1 mode (for the free end case) or the *n* = 0 mode (for the tethered end case) as the inverse of the slowest relaxation time, *τ*. That is, by diagonalizing *K* as *K* = *PλP*^*T*^ using an orthogonal matrix *P* and a diagonal matrix *λ*, we define *τ* by

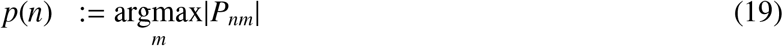

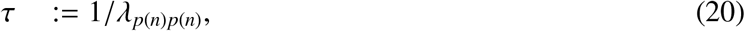

with *n* = 1for the free end chain and *n* = 0 for the tethered chain.

Under this definition, we numerically obtained *τ* from the eigenspectrum of *K*, while varying the Kuhn length *b* and fixing *N* = 96 (Extended Data Fig. 7b,c). In both the free end and tethered end cases, we obtain the crossover from the Zimm regime to the Rouse regime as *b* increases:

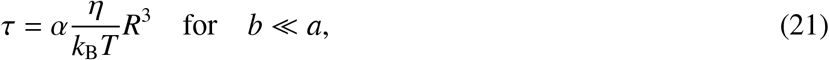

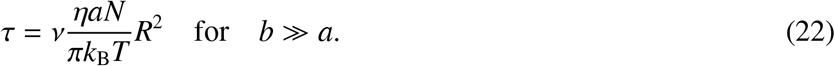

and α ≃ 1.09 for the tethered end case. We used α = 1.09 in comparing with data in Fig. 2f.

### Details for in vitro Hi-C data analysis

#### Analysis pipeline for contact map generation

The reverse complement sequence of the second read was aligned to the first read using `Bio.Align. PairwiseAligner` with the settings

- `open gap score = -2`
- `extend gap score = -4`
- `query left open gap score = -2`
- `query left extend gap score = -2`
- `query right open gap score = -2`
- `query right extend gap score = -2`
- `mode = ‘local’`

Only sequences with a score greater than 50 and the ScaI recognition sequence (AGTACT) found around the center (between 20 bp and 25 bp) were used in the downstream analysis. Groups of sequences with the same insert sequence and the UMI sequence were counted as a single sequence.

For the replicate 1 and 2 of the 96-mer experiments, we found that the non-barcoded arrays were also present at the positions of the E (NonKac) and F (Kac) arrays, likely due to misidentification of the samples during the ligation step. For these samples, we estimated the abundance of the total arrays by assuming that the contact frequency with the spike-in Kac 12-mer array is symmetric with respect to the center position of the array, and corrected the corresponding rows and columns using the estimated ratio.

20-bp sequences at the 5’ and 3’ sides of the ScaI recognition sequence were extracted, and a barcoded sequence with the minimum Hamming distance was selected. Sequences with an average Hamming distance of less than 2 (99.44% of the total sequences) were used in the downstream analysis.

#### Background level estimation

Let the count matrix *C*_*i,j*_ where *i, j* = 1, …, *M* and *i, j* = *M* + 1, …, *N* be the linker indices for the 96-mer and spike-in 12-mer arrays. We calculated the background level *B*_*i,j*_ at (*i, j*) as

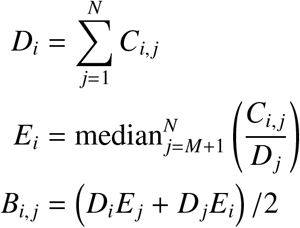

assuming 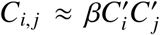 where 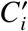 is the concentration of fragment containing the *i*-th linker and β is the efficiency for random ligation.

## Supplementary Table

**Supplementary Table S1: (Attached as a separate file) Summary of nanopore sequencing experiments**. “gDNA” protocol was used for all experiments.

## References

[1] Dixon, J. R. et al. Chromatin architecture reorgani-zation during stem cell differentiation. Nature 518, 331–336 (2015).

[2] Burton, A. & Torres-Padilla, M.-E. Chromatin dy-namics in the regulation of cell fate allocation during early embryogenesis. Nat. Rev. Mol. Cell Biol. 15, 723–734 (2014).

[3] Buenrostro, J. D. et al. Single-cell chromatin ac-cessibility reveals principles of regulatory variation. Nature 523, 486–490 (2015).

[4] Boettiger, A. N. et al. Super-resolution imaging re-veals distinct chromatin folding for different epige-netic states. Nature 529, 418–422 (2016).

[5] Xu, J. et al. Super-resolution imaging of higher-order chromatin structures at different epigenomic states in single mammalian cells. Cell Rep. 24, 873–882 (2018).

[6] Tse, C., Sera, T., Wolffe, A. P. & Hansen, J. C. Disruption of higher-order folding by core histone acetylation dramatically enhances transcription of nucleosomal arrays by RNA polymerase III. Mol. Cell. Biol. (1998).

[7] Shogren-Knaak, M. et al. Histone H4-K16 acetyla-tion controls chromatin structure and protein inter-actions. Science 311, 844–847 (2006).

[8] Allahverdi, A. et al. The effects of histone H4 tail acetylations on cation-induced chromatin fold-ing and self-association. Nucleic Acids Res. 39, 1680–1691 (2011).

[9] Funke, J. J. et al. Uncovering the forces between nucleosomes using DNA origami. Sci. Adv. 2, e1600974 (2016).

[10] Dhalluin, C. et al. Structure and ligand of a histone acetyltransferase bromodomain. Nature 399, 491–496 (1999).

[11] Lovén, J. et al. Selective inhibition of tumor onco-genes by disruption of super-enhancers. Cell 153, 320–334 (2013).

[12] Bannister, A. J. et al. Selective recognition of methylated lysine 9 on histone H3 by the HP1 chromo domain. Nature 410, 120–124 (2001).

[13] Lachner, M., O’Carroll, D., Rea, S., Mechtler, K. & Jenuwein, T. Methylation of histone H3 lysine 9 creates a binding site for HP1 proteins. Nature 410, 116–120 (2001).

[14] Machida, S. et al. Structural basis of heterochro-matin formation by human HP1. Mol. Cell 69, 385–397.e8 (2018).

[15] Poepsel, S., Kasinath, V. & Nogales, E. Cryo-EM structures of PRC2 simultaneously engaged with two functionally distinct nucleosomes. Nat. Struct. Mol. Biol. 25, 154–162 (2018).

[16] Francis, N. J., Kingston, R. E. & Woodcock, C. L. Chromatin compaction by a Polycomb group protein complex. Science 306, 1574–1577 (2004).

[17] Kilic, S. et al. Single-molecule FRET reveals multi-scale chromatin dynamics modulated by HP1α. Nat. Commun. 9, 235 (2018).

[18] Song, F. et al. Cryo-EM study of the chromatin fiber reveals a double helix twisted by tetranucleosomal units. Science 344, 376–380 (2014).

[19] Gibson, B. A. et al. Organization of chromatin by intrinsic and regulated phase separation. Cell 179, 470–484 (2019).

[20] Strickfaden, H. et al. Condensed chromatin behaves like a solid on the mesoscale in vitro and in living cells. Cell 183, 1772–1784.e13 (2020).

[21] Portillo-Ledesma, S., Li, Z. & Schlick, T. Genome modeling: from chromatin fibers to genes. Curr. Opin. Struct. Biol. 78, 102506 (2023).

[22] Mishra, L. N. & Hayes, J. J. A nucleosome-free region locally abrogates histone H1–dependent re-striction of linker DNA accessibility in chromatin. J. Biol. Chem. 293, 19191–19200 (2018).

[23] Müller, M. M., Fierz, B., Bittova, L., Liszczak, G. & Muir, T. W. A two-state activation mechanism con-trols the histone methyltransferase Suv39h1. Nat. Chem. Biol. 12, 188–193 (2016).

[24] Blacketer, M. J., Feely, S. J. & Shogren-Knaak, M. A. Nucleosome interactions and stability in an ordered nucleosome array model system. J. Biol. Chem. 285, 34597–34607 (2010).

[25] Lowary, P. T. & Widom, J. New DNA sequence rules for high affinity binding to histone octamer and sequence-directed nucleosome positioning. J. Mol. Biol. 276, 19–42 (1998).

[26] Turner, B. M. Histone acetylation and control of gene expression. J. Cell Sci. 99, 13–20 (1991).

[27] Grunstein, M. Histone acetylation in chromatin structure and transcription. Nature 389, 349–352 (1997).

[28] Wakamori, M. et al. Intra-and inter-nucleosomal interactions of the histone H4 tail revealed with a human nucleosome core particle with genetically-incorporated H4 tetra-acetylation. Sci. Rep. 5, 17204 (2015).

[29] Wakamori, M. et al. Quantification of the effect of site-specific histone acetylation on chromatin tran-scription rate. Nucleic Acids Res. 48, 12648–12659 (2020).

[30] Shipony, Z. et al. Long-range single-molecule map-ping of chromatin accessibility in eukaryotes. Nat. Methods 17, 319–327 (2020).

[31] Stergachis, A. B., Debo, B. M., Haugen, E., Church-man, L. S. & Stamatoyannopoulos, J. A. Single-molecule regulatory architectures captured by chromatin fiber sequencing. Science 368, 1449–1454 (2020).

[32] Chen, Q., Yang, R., Korolev, N., Liu, C. F. & Nor-denskiöld, L. Regulation of nucleosome stacking and chromatin compaction by the histone H4 N-terminal tail–H2A acidic patch interaction. J. Mol. Biol. 429, 2075–2092 (2017).

[33] Zimm, B. H. Dynamics of polymer molecules in di-lute solution: viscoelasticity, flow birefringence and dielectric loss. J. Chem. Phys. 24, 269–278 (1956).

[34] Doi, M. & Edwards, S. F. The theory of polymer dynamics, vol. 73 (Oxford University Press, 1988).

[35] Yao, J., Lowary, P. T. & Widom, J. Direct detection of linker DNA bending in defined-length oligomers of chromatin. Proc. Natl. Acad. Sci. 87, 7603–7607 (1990).

[36] Goldman, A. J., Cox, R. G. & Brenner, H. Slow vis-cous motion of a sphere parallel to a plane wall—i motion through a quiescent fluid. Chem. Eng. Sci. 22, 637–651 (1967).

[37] Lieberman-Aiden, E. et al. Comprehensive map-ping of long-range interactions reveals folding prin-ciples of the human genome. Science 326, 289–293 (2009).

[38] Oberbeckmann, E., Quililan, K., Cramer, P. & Oudelaar, A. M. In vitro reconstitution of chromatin domains shows a role for nucleosome positioning in 3D genome organization. Nat. Genet. 56, 483–492 (2024).

[39] Ohno, M. et al. Sub-nucleosomal genome structure reveals distinct nucleosome folding motifs. Cell 176, 520–534 (2019).

[40] Maeshima, K. et al. Nucleosomal arrays self-assemble into supramolecular globular structures lacking 30-nm fibers. EMBO J. 35, 1115–1132 (2016).

[41] Olins, D. E. & Olins, A. L. Chromatin history: our view from the bridge. Nat. Rev. Mol. Cell Biol. 4, 809–814 (2003).

[42] Routh, A., Sandin, S. & Rhodes, D. Nucleosome repeat length and linker histone stoichiometry de-termine chromatin fiber structure. Proc. Natl. Acad. Sci. 105, 8872–8877 (2008).

[43] Jentink, N., Purnell, C., Kable, B., Swulius, M. T. & Grigoryev, S. A. Cryoelectron tomography reveals the multiplex anatomy of condensed native chromatin and its unfolding by histone citrullina-tion. Mol. Cell 83, 3236–3252 (2023).

[44] Cai, S., Song, Y., Chen, C., Shi, J. & Gan, L. Natural chromatin is heterogeneous and self-associates in vitro. Mol. Biol. Cell 29, 1652–1663 (2018).

[45] Crane, E. et al. Condensin-driven remodelling of X chromosome topology during dosage compensation. Nature 523, 240–244 (2015).

[46] Halverson, J. D., Smrek, J., Kremer, K. & Grosberg, A. Y. From a melt of rings to chromosome territo-ries: The role of topological constraints in genome folding. Rep. Prog. Phys. 77, 022601 (2014).

[47] Sanborn, A. L. et al. Chromatin extrusion explains key features of loop and domain formation in wild-type and engineered genomes. Proc. Natl. Acad. Sci. 112, E6456–E6465 (2015).

[48] Adachi, K. & Kawaguchi, K. Chromatin state switching in a polymer model with mark-conformation coupling. Phys. Rev. E 100, 060401 (2019).

[49] Barbieri, M. et al. Complexity of chromatin folding is captured by the strings and binders switch model. Proc. Natl. Acad. Sci. 109, 16173–16178 (2012).

[50] Collepardo-Guevara, R. et al. Chromatin unfolding by epigenetic modifications explained by dramatic impairment of internucleosome interactions: a mul-tiscale computational study. J. Am. Chem. Soc. 137, 10205–10215 (2015).

[51] Michieletto, D., Orlandini, E. & Marenduzzo, D. Polymer model with epigenetic recoloring reveals a pathway for the de novo establishment and 3D or-ganization of chromatin domains. Phys. Rev. X 6, 041047 (2016).

## References

[1] Yao, J., Lowary, P. T. & Widom, J. Direct detection of linker DNA bending in defined-length oligomers of chromatin. Proc. Natl. Acad. Sci. 87, 7603–7607 (1990).

[2] Lowary, P. T. & Widom, J. New DNA sequence rules for high affinity binding to histone octamer and sequence-directed nucleosome positioning. J. Mol. Biol. 276, 19–42 (1998).

[3] Kilic, S. et al. Single-molecule FRET reveals multiscale chromatin dynamics modulated by HP1α. Nat. Commun. 9, 235 (2018).

[4] Kikuchi, M. et al. Epigenetic mechanisms to propagate histone acetylation by P300/CBP. Nat. Com-mun. 14, 4103 (2023).

[5] Mivelaz, M. & Fierz, B. Observing protein interaction dynamics to chemically defined chromatin fibers by colocalization single-molecule fluorescence microscopy. Methods 184, 112–124 (2020).

[6] Tokunaga, M., Imamoto, N. & Sakata-Sogawa, K. Highly inclined thin illumination enables clear single-molecule imaging in cells. Nat. Methods 5, 159–161 (2008).

[7] Edelstein, A., Amodaj, N., Hoover, K., Vale, R. & Stuurman, N. Computer control of microscopes using µManager. Curr. Protoc. Mol. Biol. 92, 14.20.1–14.20.17 (2010).

[8] Pinkard, H. et al. Pycro-Manager: Open-source software for customized and reproducible microscope control. Nat. Methods 18, 226–228 (2021).

[9] Ershov, D. et al. TrackMate 7: Integrating state-of-the-art segmentation algorithms into tracking pipelines. Nat. Methods 19, 829–832 (2022).

[10] Tinevez, J.-Y. et al. TrackMate: An open and extensible platform for single-particle tracking. Methods 115, 80–90 (2017).

[11] Vestergaard, C. L., Blainey, P. C. & Flyvbjerg, H. Optimal estimation of diffusion coefficients from single-particle trajectories. Phys. Rev. E 89, 022726 (2014).

[12] Oberbeckmann, E., Quililan, K., Cramer, P. & Oudelaar, A. M. In vitro reconstitution of chromatin domains shows a role for nucleosome positioning in 3D genome organization. Nat. Genet. 56, 483–492 (2024).

[13] Oxford Nanopore Technologies PLC. Nanoporetech/dorado. URL https://github.com/nanoporetech/dorado.

[14] Oxford Nanopore Technologies PLC. Nanoporetech/modkit. URL https://github.com/nanoporetech/modkit.

[15] van der Walt, S. et al. Scikit-image: Image processing in Python. PeerJ 2, e453 (2014).

[16] Knight, P. A. & Ruiz, D. A fast algorithm for matrix balancing. IMA J. Numer. Anal. 33, 1029–1047 (2013).

## References

[1] Dorigo, B., Schalch, T., Bystricky, K. & Richmond, T. J. Chromatin fiber folding: Requirement for the histone H4 N-terminal Tail. J. Mol. Biol. 327, 85–96 (2003).

[2] Kilic, S. et al. Single-molecule FRET reveals multiscale chromatin dynamics modulated by HP1α. Nat. Commun. 9, 235 (2018).

[3] Zhang, Z., Park, S., Pertsinidis, A. & Revyakin, A. Cloud-point PEG glass surfaces for imaging of immobilized single molecules by total-internal-reflection microscopy. BIO-PROTOCOL 6 (2016).

[4] Tokunaga, M., Imamoto, N. & Sakata-Sogawa, K. Highly inclined thin illumination enables clear single-molecule imaging in cells. Nat. Methods 5, 159–161 (2008).

[5] Rohland, N. & Reich, D. Cost-effective, high-throughput DNA sequencing libraries for multiplexed target capture. Genome Res. 22, 939–946 (2012).

[6] Nečas, D. & Klapetek, P. Gwyddion: an open-source software for SPM data analysis. Cent. Eur. J. Phys. 10, 181–188 (2012).

[7] Kubo, S., Umeda, K., Kodera, N. & Takada, S. Removing the parachuting artifact using two-way scanning data in high-speed atomic force microscopy. Biophys. Physicobiol. 20 (2023).

[8] Erb, D. pybaselines: A Python library of algorithms for the baseline correction of experimental data. URL https://github.com/derb12/pybaselines.

[9] Cock, P. J. A. et al. Biopython: Freely available Python tools for computational molecular biology and bioinformatics. Bioinformatics 25, 1422–1423 (2009).

[10] van der Walt, S. et al. Scikit-image: Image processing in Python. PeerJ 2, e453 (2014).

[11] Roelfs, M. & Kroon, P. C. tBuLi/symfit: Symfit 0.5.6. Zenodo (2023). URL 10.5281/zenodo.7643591.

[12] Fukai, Y. T. & Kawaguchi, K. LapTrack: Linear assignment particle tracking with tunable metrics. Bioinformatics 39, btac799 (2023).

[13] Koch, M., Sommer, J.-U. & Blumen, A. Polymer chains tethered to impenetrable interfaces: Broadening of relaxation spectra. J. Chem. Phys. 106, 1248–1256 (1997).

